# Clonally heritable gene expression imparts a layer of diversity within cell types

**DOI:** 10.1101/2022.02.14.480352

**Authors:** Jeff E. Mold, Martin H. Weissman, Michael Ratz, Michael Hagemann-Jensen, Joanna Hård, Carl-Johan Eriksson, Hosein Toosi, Joseph Berghenstråhle, Leonie von Berlin, Marcel Martin, Kim Blom, Jens Lagergren, Joakim Lundeberg, Rickard Sandberg, Jakob Michaëlsson, Jonas Frisén

## Abstract

Cell types can be classified based on shared patterns of transcription. Variability in gene expression between individual cells of the same type has been ascribed to stochastic transcriptional bursting and transient cell states. We asked whether long-term, heritable differences in transcription can impart diversity within a cell type. Studying clonal human lymphocytes and mouse brain cells, we uncover a vast diversity of heritable transcriptional states among different clones of cells of the same type in vivo. In lymphocytes we show that this diversity is coupled to clone specific chromatin accessibility, resulting in distinct expression of genes by different clones. Our findings identify a source of cellular diversity, which may have important implications for how cellular populations are shaped by selective processes in development, aging and disease.

## Main

Multicellular organisms are composed of diverse cell types, which can be classified according to shared patterns of gene expression (*1, 2*). Transcriptome-wide single cell profiling separates cells into distinct cell types, and at the same time reveals transcriptional variability among cells of a single type (*3*). Variability in gene expression within cell types is thought to reflect stochastic transcription or transient fluctuations of phenotype, often referred to as cell states (*4*). We asked whether an additional mechanism-heritable, clonal differences - may contribute to variability observed within a cell type. Such differences would impart unique clonotypic features upon the progeny of individual cells, which could explain some of the diversity seen within cell types and have implications for cell selection in health and disease (*5*). Evidence exists for short-term heritability of gene expression states in transformed cell lines in vitro (*6–9*), but this has not been much explored in primary cells or in vivo at a transcriptome-wide level. Here we assessed whether stable clonally heritable gene expression programs contribute to diversity in long-lived human lymphocyte subsets as well as in cells of the mouse central nervous system in vivo.

### Expanded T cell clones show evidence of heritable clonal gene expression in vivo

We took advantage of the genetic barcodes arising from T cell receptor (TCR) rearrangement to study clonally expanded populations of cells that develop from individual naïve CD8+ T cells in humans after vaccination with yellow fever virus vaccine (YFV-17D). We analyzed the transcriptomes of 3,837 HLA-A2/YFV-specific CD8+ T cells from three healthy donors using high-sensitivity, full transcript single cell RNA-seq (Smart-seq3) (table S1) (*10*). We identified single cells belonging to expanded clonal populations in the circulating blood during the memory phase of the immune response (Donors A, B: Day 180, Donor C: Day 1,286 post-vaccination) according to shared TCR sequences (*11*). To analyze clonal gene expression differences, we selected the 10 largest clones from each donor (205, 252, 203 cells in total in donors A, B, C respectively, with at least 14 cells/clone) and identified differentially expressed genes between clones in each donor independently using an ANOVA F-test (unadjusted p < 0.05, Fig. 1A, Fig. S1A, table S2). We computed the same statistics using 1000 permutations of clone labels to estimate the number of false discoveries. Using the 95^th^ percentile among these 1000 permutations as a conservative estimate of false discoveries, we found 175, 268, and 323 genes with interclonal differential expression (interclonal variability), in excess beyond false discoveries, in donors A, B and C, respectively (**Fig. 1A**, methods).

**Figure 1.**
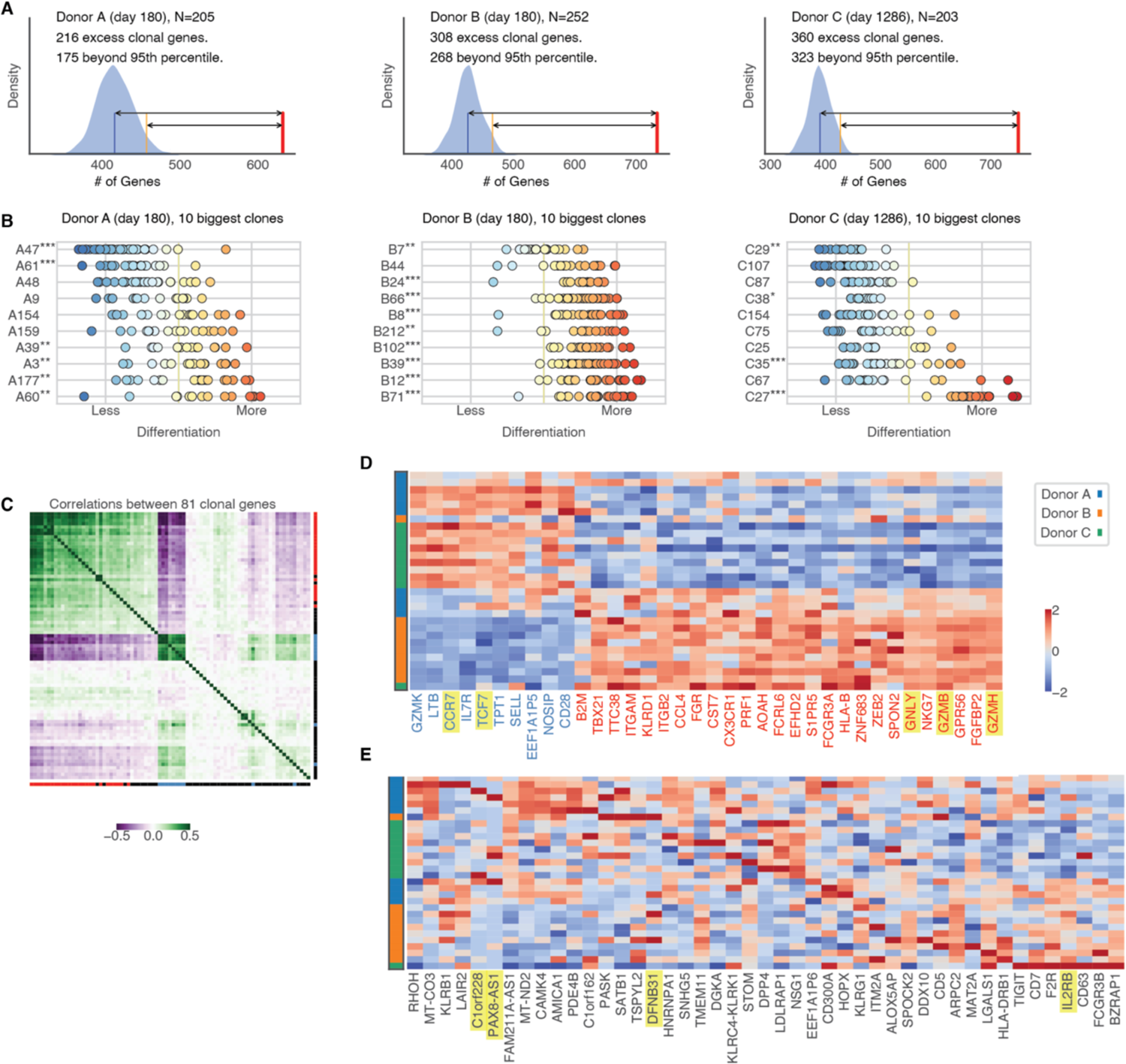
Heritable Transcriptional States in Expanded Clonal T cells In Vivo. **(A)** Numbers of clonally variable genes found in top 10 largest clones for Donors A-C based on ANOVA F-statistic. Timepoint post-vaccination and cell numbers are shown near donor ID. Here, clonally variable genes are those with unadjusted p<0.05, and their number is estimated by comparison to the 95th percentile among 1000 permutations of clone labels (blue KDE-smoothed histogram). **(B)** Distribution of cells from each clone and donor, according to differentiation state based on PC1 from clonally variable genes. Clones showing strong bias as compared to the full donor population are labeled (one-sample Kolmogorov-Smirnov test, two-sided p-value, *<0.05, **<0.01. ***<0.001). **(C)** Correlation plot indicating highly correlated (green) and anti-correlated (purple) modules among the most clonally variable genes (excluding RPL/RPS genes). Genes are marked with red/blue if they are associated with differentiation state, and black otherwise. **(D,E)** Heatmaps indicating average clonal expression levels of genes with high levels of contribution to PC1 (**D**) and other clonal genes not associated with differentiation states (**E**). Clones are ordered according to average PC1 score of all individual cells. Donors are indicated by color (Blue: A, Orange: B, Green: C) and rows depict average gene expression level of each clone (z-score).

Principal component analysis performed on the top interclonally differentially expressed genes (106 genes, estimated FDR < 3%) identified a subset of genes linked to T cell differentiation states (**fig S1, A-D**) (*11–13*). These genes contributed strongly to PC1 and spread the CD8+ T cells from all donors along the established continuum of differentiation states observed in memory CD8+ T cell populations (**Fig. 1B**) (*11, 12, 14*). Clonally related cells exhibited biases along this continuum of differentiation states, which persisted for years after the initial activation and expansion phase of the response.

On the other hand, 45 of the most interclonally differentially expressed genes did not contribute strongly to PC1. A correlation analysis showed neither significant correlations between these genes and those associated with differentiation state, nor correlations among these 45 genes (**Fig. 1, C-E**). Some of these genes were expressed sporadically in the population, but often by many cells from individual clones (*PAX8-AS1*, *C1orf228*, *DNFB31, PASK, SATB1*) (**Fig. 1E**). Others were expressed frequently among all cells yet exhibited variable expression ranges between different clonal populations (*IL2RB*, *CD7*).

We also observed that some genes associated with differentiation state exhibited interclonal differential expression even among clones with similar differentiation bias (*ZNF683, GNLY, GZMB, GZMH*), suggesting complex patterns of transcriptional differences even within highly correlated gene modules (**Fig. 1E**). Thus, we identify two layers of heritable clonal transcriptional traits in long-lived memory T cell clones: (1) differentiation biases involving highly correlated blocks of genes frequently expressed across all memory CD8+ T cells and (2) gene expression differences which appears sporadic at the population level but which are restricted to a more narrow range within individual clones.

### Unique clonal transcriptional states emerge upon reactivation of individual T cells

Long-lived memory T cells exist in a resting state and may circulate throughout the body for years between cell divisions (*15*). Reactivation of memory T cells leads to rapid proliferation and differentiation of this population, revealing a complex pattern of transcriptional activity not observed in the resting state (*13, 16*). Therefore, we decided to investigate clonally heritable transcriptional profiles after activation and differentiation of memory T cells in short-term in vitro cultures using a previously generated dataset (*17*).

We examined nine distinct T cell clones (31-48 cells sampled from each clone), each expanded from a single memory T cell isolated 136 days post-vaccination (Donor D) (**Fig. 2A**). Based on total cell numbers after expansion, each clone was estimated to have undergone 10-12 rounds of division during 19 days in cell culture. High-coverage, full-transcript single cell transcriptomes were generated using Smart-seq2, and gene counts were normalized based on transcripts per million reads (TPM) (*18, 19*). After filtering genes by discarding all TCR genes and removing lowly expressed genes, 7,440 genes remained for analysis. We measured interclonal differential gene expression among all genes with ANOVA (parametric) and Kruskal-Wallis (non-parametric) tests and identified 2,034 genes which varied significantly between populations of clonally related cells (unadjusted p<0.01) in either test (1,488 in both tests, table S3). In contrast, only 98 genes met this threshold for significance (p<0.01) when performing the tests with a random permutation of clone labels.

**Figure 2.**
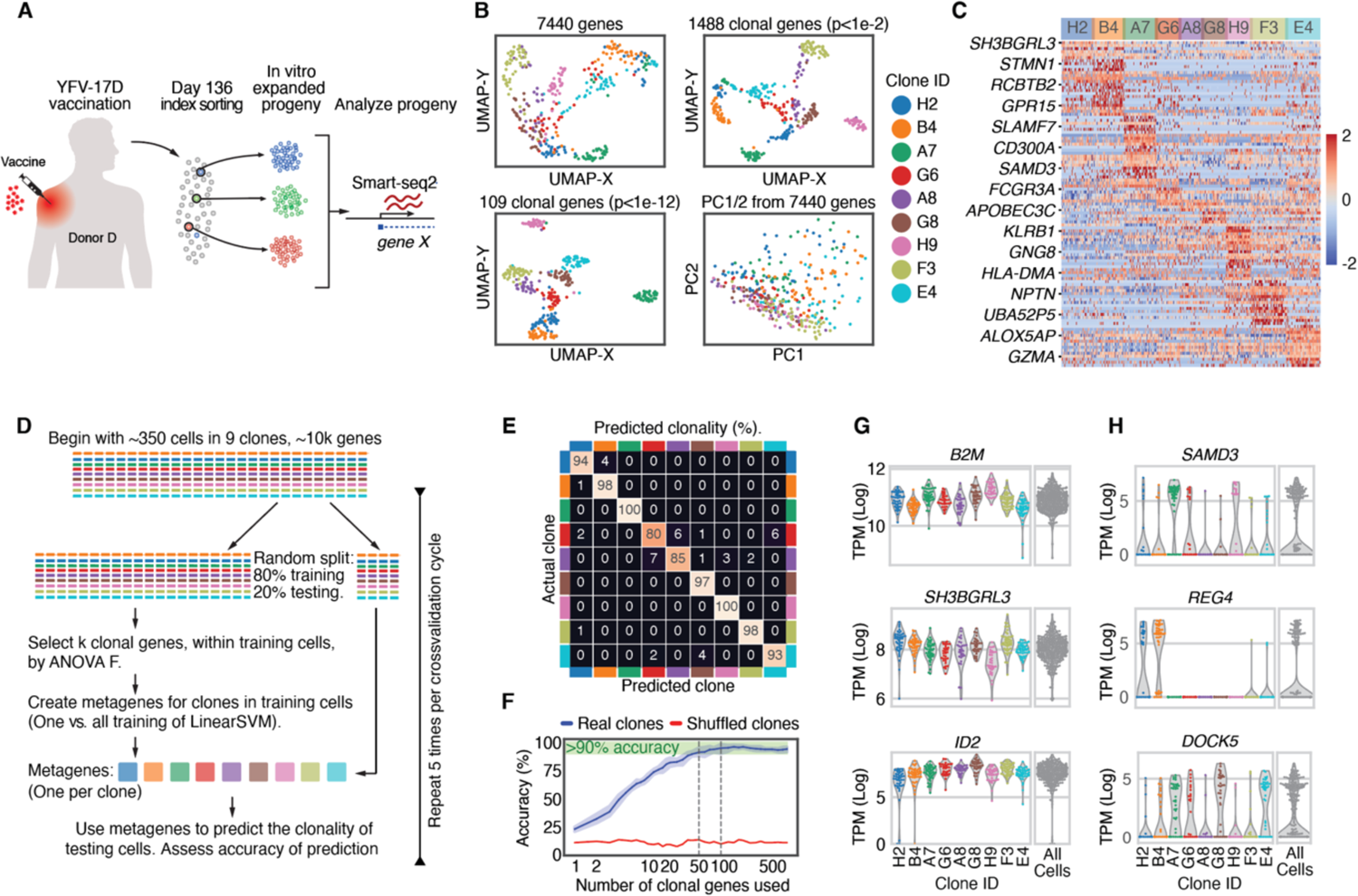
Heritable transcriptional states in expanded clonal T cells in vitro. **(A)** Schematic illustrating experimental strategy for isolating and expanding individual T cells in vitro. Single HLA-A2/YFV NS4b-specific memory CD8+ T cells were index-sorted and expanded with irradiated autologous feeder cells, IL-2 and NS4b peptide for 21 days prior to analyses. **(B)** Visualization of 352 cells from 9 different clones based on UMAP analysis using the ten first principal components (PCs) by PCA on all genes (7,440 genes, excluding TCR components and low expressed genes) (top left square), on clonally variable genes defined by both ANOVA F-statistic and Kruskal-Wallis as significant (n=1,488 genes; p<1e-2, top right) or highly significant (n=109 genes; p<1e-12, bottom left). The clonal distribution based on the two first PCs is shown in lower right plot. **(C)** Heatmap displaying 109 clonally variable genes defining distinct clonal transcriptome profiles. **(D)** Schematic illustrating strategy to test identification of single cells based on SVM classifier. **(E)** Confusion matrix displaying accuracy for each clonal population. **(F)** Prediction accuracy for all clones (real clones) compared to prediction accuracy of test performed on randomly assigned ‘clones’ (shuffled clones) relative to numbers of genes included for prediction. **(G)** Examples of highly expressed genes which show clonal variability (‘tunable’ genes) and **(H)** genes which are either ON or OFF in the majority of cells from each clone. All 352 cells are shown together in gray on right hand of each plot.

Some clonal structure was apparent when reducing the dimensionality from all 7,440 genes to the ten first dimensions (PC1-10) by PCA followed by UMAP visualization (**Fig. 2B**), and nearly unambiguous clustering of clonally related cells was achieved when performing the same analysis using highly significant clonal genes (**Fig. 2B**) (*20*). In contrast, using only the top two principal components (PC1-2) based on all 7,440 genes revealed no clear clonal structure in the data, suggesting that heritable clonal gene expression patterns cannot be explained by a few coordinated blocks of genes (**Fig. 2B**). Clonal gene expression differences were also clear when visualizing patterns of expression for genes with significant interclonal differential expression (**Fig. 2C**).

To assess whether the clonal identities of individual T cells could be determined from single cell gene expression signatures, without TCR information, we applied a machine learning classifier (linear support vector machine, SVM). For cross-validation, we trained the SVM classifier on 80% of cells from each clone to select clonal genes and create 9 metagenes (SVM hyperplanes). These metagenes were used to predict the clonal origins of the remaining 20% of cells (**Fig. 2D**). This approach placed individual cells in their respective clones, with accuracy ranging from 80-100% for each clone (**Fig. 2, E and F**), estimated by 100 repetitions of the training/testing procedure with 100 or more clonally variable genes selected each time. Similar levels of accuracy were achieved when using 50% of the cells in each clone for training.

Performing the same analysis with randomly permuted clone labels gave no predictive power beyond chance (**Fig. 2F**). We observed that accuracy of the prediction increased with the number of selected genes, reaching 95% accuracy with 100 or more genes, further indicating that clones are not identified simply by a small number of rarely expressed genes (**Fig. 2F**). Closer examination of genes with significant interclonal differential expression revealed both genes expressed at a high level throughout the population, but with distinct clonally variable ranges of expression (e.g. *B2M*, *SH3BGRL3*, *ID2*), as well as genes expressed only in certain clones (e.g. *SAMD3*, *REG4*, *DOCK5*) (**Fig. 2, G and H**). Some highly expressed interclonally differentially expressed genes were previously identified as heritably maintained in short-term cultures of transformed cell lines and in cancer clones in vivo (*8, 21*).

We next analyzed three highly expanded clonal populations (16-17 doublings over 20 days) from an additional donor isolated at day 2,001 post-vaccination (Donor E). We profiled 598 single cells (clone A: 277, clone B: 162, clone C: 154) using Smart-seq3 enabling direct comparison between UMI and TPM-based normalization of gene expression values. We detected comparable numbers of interclonally differentially expressed genes whether using UMI or TPM-normalized datasets and no clear relationships between interclonal variability and transcript expression levels, cell size or granularity (**Fig. S2, A-C, table S4**). Interestingly, with larger numbers of cells and fewer clones, PCA clearly separated single cells into the three clones A, B, and C, just by considering PC1 and PC2 (**Fig. S2D**). Furthermore, clone A visibly split into two subgroups of cells, which were made precise by Louvain clustering. The two subgroups appeared to have undergone distinct differentiation trajectories during activation and expansion based on phenotyping of surface marker expression by flow cytometry (**Fig. S2E**). This was further demonstrated by examining a large set of genes enriched within each clonal population, which revealed shared clonal features of all cells within clone A as well as evidence of subclonal diversification (**Fig. S2, F and G**). These findings confirm the diversity of interclonal variability in gene expression, and its heritability even in the presence of subclonal diversification.

### Stable maintenance of clonal transcriptional states in memory T cells for over a year in vivo

Late in the memory phase of the immune response, the clonal diversity of the circulating memory T cell pool decreases, increasing the likelihood that independently sorted T cells share a clonal origin, and we refer to such cells as sisters (*11*). Because circulating memory T cells continuously migrate between distinct lymphoid tissues, and possibly other peripheral organs/tissues, it is likely that these cells have experienced substantially different environmental exposures over the course of their individual lifespans (*22*). Nonetheless we observed that clonally related cells in vivo often shared heritable patterns of transcription (**Fig. 1**). To address whether resting clonally related memory T cells in vivo produced progeny with similar clonal transcriptional profiles after activation and differentiation in vitro, we generated a dataset with 24 expanded clones isolated late in the memory phase of the response (day 593 post vaccination) and identified four sets of sister clones (**Fig. 3A**). Cross-validated linear SVM classification on all 20 clones (combining sister clones from different wells) once again gave a high degree of accuracy of clonal identification (88% on average) for single cells from all 20 clones, indicating that each clone possessed a distinct heritable transcriptional signature (**Fig. 3B, Table S5**).

**Figure 3.**
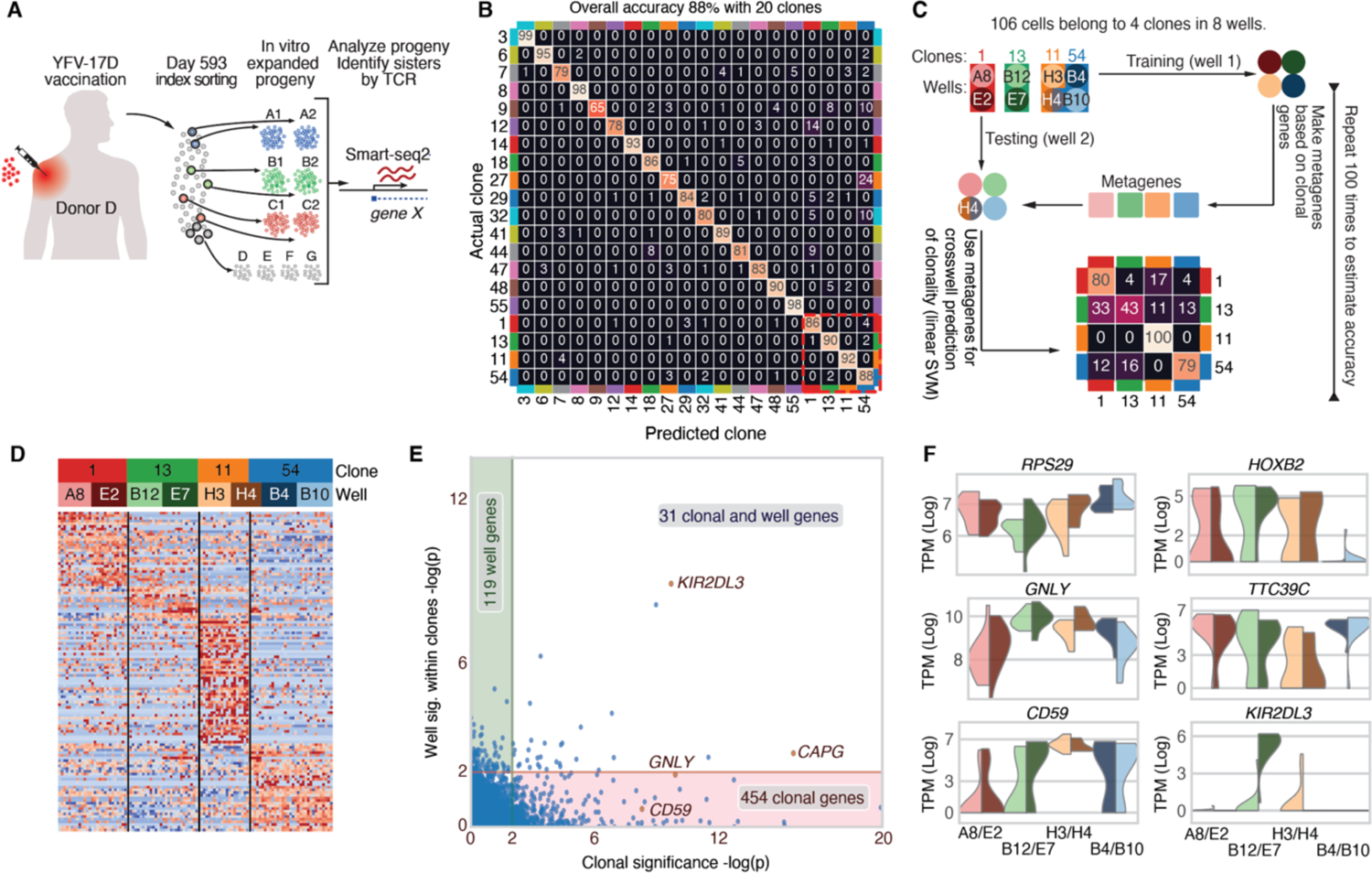
Shared transcriptional identities of progeny from ‘sister’ clones separated in vivo. **(A)** Schematic illustrating experimental strategy for isolating and expanding sister clones in vitro. **(B)** Confusion matrix showing SVM prediction accuracies for all clonal cells (defined by TCR sequence, combining sisters in separate wells (clones 1, 13, 11 and 54)). **(C)** Schematic illustrating strategy to measure prediction accuracy by SVM algorithm trained on clonal T cells from one sister (in a single well) to predict cells from the second sister (in a separate well). One well (H4) contained a mixture of two clones (cl11 and cl54) and was used as a test well. Prediction accuracies are reported in confusion matrix shown underneath schematic illustration. **(D)** Heatmap showing clonal genes which were shared by clonally related cells derived from each sister in separate wells. Well H4 is bisected to indicate cells from clones 11 and 54 respectively. **(E)** Nested ANOVA test to estimate the number of genes which show significant variability according to which well they are from versus which clonal origin they share. Genes on the Y-axis (well genes: 119) show significant transcriptional differences arising during activation in a given well independent of clonal relationships between wells. X-axis genes (clonal genes: 454) are clonally variable and shared by cells from distinct sisters. **(F)** Split violin plots showing clonally variable gene expression patterns by sister clones. *KIR2DL3* is a rare example of a clonally variable gene which also varies between sisters.

Focusing on sister clones, we determined whether the progeny of one sister (one well) exhibited transcriptional profiles similar to the progeny from the other sister. Here we trained an SVM classifier on all cells from one well (sisters A) for each of the four sets of sister clones, and used the identified clonal signatures to predict the clonal identity of cells from the remaining wells (sisters B, **Fig. 3C**). For 3 clones (Clones 1, 11, and 54) we observed 80, 100, and 79% prediction accuracy respectively. The fourth clone (clone 13) was predicted with lower accuracy (43%), reflecting more substantial differences between the wells containing each expanded sister clone, and fewer genes separating this clone from the other three (**Fig. 3D**). In the broader context of all 20 clones, single cells from clone 13 (pooling both wells) were classified with high accuracy (90%) (Fig. 3b). This suggests that the clone had diversified somewhat between the two sisters yet retained a clonal transcriptional signature.

In addition to genes with shared patterns of expression between sister clones, we also observed genes which varied between them. Some of the strongest differences observed between sister clones included genes which are known to exhibit fixed clonal expression patterns by lymphocytes throughout their maturation (e.g. *KIR* genes) (*23*). Using a nested ANOVA test, we identified 454 genes with significant (p<0.01) interclonal differential gene expression, but minimal intraclonal variability between sisters in separate wells (**Fig. 3E, table S5**). Conversely, we found 119 genes which showed significant (p<0.01) intraclonal variability between sisters, but whose interclonal variability was insignificant (beyond that which could be explained by well differences). Finally, 31 genes exhibited both inter- and intraclonal variability. This is consistent with our prior assessment that subclonal diversification can arise within clonally related cells during cell division, while an overarching clonal signature remains.

By using TCR sequences to confirm the clonal identities of all single cells in each well, we noted that one well contained an admixture of cells from two different clones, likely due to technical errors while sorting the founder cells for these clonal expansions (accounted for in **Fig. 3, B-F**). Well H4 was found to contain cells from clones 11 and 54 in roughly equal numbers, indicating that memory T cells corresponding to each clone were sorted together into this well. Despite this admixture occurring, cells from well H4 retained gene expression patterns similar to their sisters in other wells (clone 11 sisters in well H3, and clone 54 sisters in wells B10/B4) (**Fig. 3, C and D**). This was particularly apparent when comparing clone-specific gene expression patterns between clones 11 and 54 side by side (**Fig. 3D**). This fortuitous experiment demonstrates that diverse interactions between unrelated cells within an enclosed environment was not a major contributor to clonal gene expression differences.

### Heritable phenotypes are not due to genomic copy number variations

Because genetic mutations can occur during somatic cell division, we asked whether clones experienced extensive genomic copy number variations (CNV) during somatic clonal expansions. We performed single cell CNV analysis on 4 single cells from each of the 24 clonal expansions in the previous dataset (**Fig. S3, A and B**) (*24*). We observed no consistent evidence of CNV (500kbp bins) in the genomes of each clone in our set of 24 expanded clones (Fig. 3), with the exception of one clone exhibiting loss of Y chromosome, known to spontaneously occur in human lymphocytes (*25*). In this clone (clone 12) we observed a loss of Y-chromosome gene expression as well as a 50% reduction in the average expression of *CD99* which is a pseudoautosomal gene expressed on both X and Y chromosomes (**Fig. S3B**).

### Clonal gene expression differences impact protein expression levels

We collected information about cell surface expression of several proteins on the progeny of all 24 expanded T cells from the previous experiment (identifying sisters separately). This allowed us to address whether differences in mRNA expression levels translated to differences in protein levels in single cells and between different clones (table S6). Protein levels correlated poorly with mRNA expression across the entire population of CD8+ T cells in our dataset (coefficient of determination range r^2^=0.001-0.190), consistent with the understanding that noise in either measurement and that transcription often occurs in bursts results in weak mRNA-protein relationships in single cells surveyed at a snapshot in time (*26*). We observed a substantially stronger correlation when analyzing the average mRNA and protein expression levels for each clone (range r^2^=0.018-0.556, **Fig. S4**). This was true for highly expressed proteins which define cell type (CD8A) as well as variably expressed proteins reflecting different activation or differentiation states (CD95/FAS, CD27, PD-1). Working with clonal averages, the highly non-normal distribution of mRNA abundance at the single-cell level is replaced by a nearly normal distribution around the clonal mean. These findings indicated that clones exhibit variable set points in mRNA abundance, which give rise to downstream variabilities in protein expression.

### Variable chromatin accessibility linked to clonally distinct transcriptional profiles

Gene expression is determined by transcription factor activity on proximal promoters and regulatory regions of DNA, collectively referred to as cis regulatory elements (CREs) (*27*). Identifying accessible regions of chromatin with ATAC-seq has become a fundamental tool for assessing potential epigenetic heterogeneity between populations of cells, revealing hidden layers of gene regulation across cell types and differentiation states (*28, 29*). Because we observed complex patterns of heritable gene expression in T cell clones, we sought an explanation in clonally variable patterns of CRE accessibility. For this goal, we generated 23 clonal expansions of T cells taken 1,401 days post-vaccination, splitting the expanded clonal populations to perform bulk ATAC-seq (1-2 replicates) and RNA-seq (3 replicates) on each (**Fig. 4A**) (*30*). For six clones, we created 2-replicate ATAC-seq samples, which demonstrated a high degree of similarity, indicating little technical noise (**Fig. 4B**). We assigned a clonal variability score (Relative Peak Variability, RPV) to each ATAC-seq peak by comparing the range of peak heights among all clones to the range expected from technical noise (see methods, **Fig. S5, A-D** for detailed explanation, table S7). Using this interclonal difference metric, we identified 9,846 significant interclonally different peaks out of 26,040 high confidence CREs detected in our clonal dataset (RPV>1, peak height range among clones greater than maximum expected from technical noise). This represents 37.8% of all CREs detected using stringent filters to remove potentially noisy peaks (normalized peak heights <30 for all clones). These variable peaks were found to be evenly distributed among promoter and putative enhancer regions (**Fig. 4C**). Interestingly we found an enrichment of interclonally variable peaks (RPV>1.5) near interclonally differentially expressed genes from all datasets. To assess this enrichment, we compared interclonally differentially expressed genes to control gene sets developed from random permutations of clone labels (**Fig. S5, E and F**). This enrichment was present, even for a large set of interclonally differentially expressed genes in vivo where we estimate a much higher false discovery rate (**Fig. 1A, Fig. S5F, table S2**).

**Figure 4.**
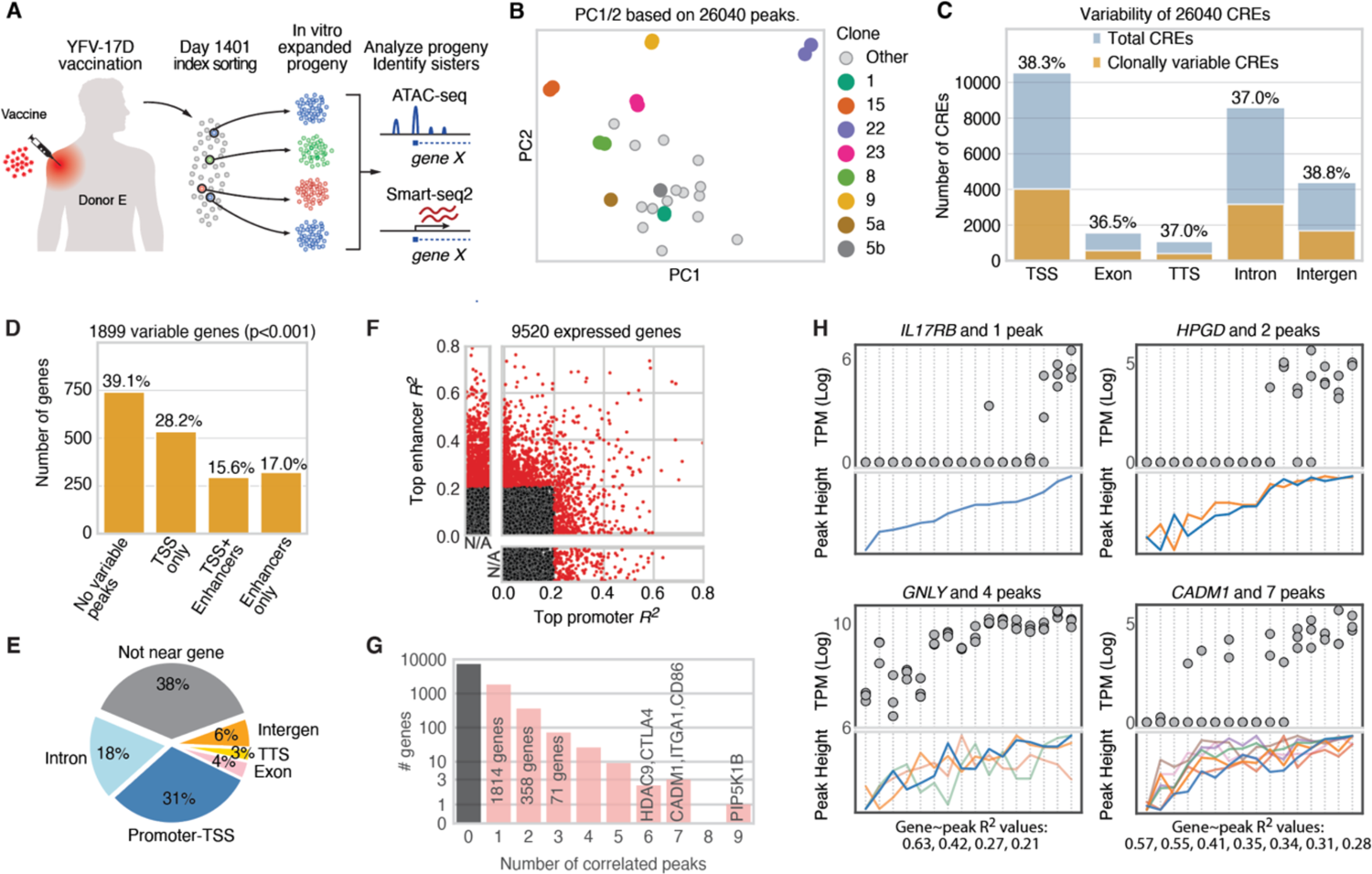
Heritable differences in chromatin accessibility underlie clonally variable gene expression. **(A)** Experimental approach to collect matched RNA and ATAC-seq datasets from clonally expanded T cells. **(B)** PCA performed on all clonal samples using 26,040 high quality peaks. Biological replicates cluster tightly together (colored dots) and two sister clones (5a and 5b) are separated in PC1. **(C)** Fraction of peaks showing evidence of clonal variability (RPV>1) in genomic positions annotated as promoter (TSS) and enhancer regions. The 26,040 CREs analyzed reach peak height >=30 for at least one clone. **(D)** Relationship between variable CRE locations and genes which show clonally variable expression patterns in matched RNA-seq analysis. **(E)** Distribution of 9,846 clonally variable CREs (RPV>1). CREs not located within 50kbp of an expressed gene in matched RNA-seq dataset (in any clone) are considered ‘not near gene’. **(F)** Scatterplot showing correlation between peak heights and gene expression for all expressed genes (9,520 genes) in the RNA-seq dataset. 16 clonal populations are included in this analysis (including 5a and 5b). Red dots indicate CRE-gene relationships with R^2^ >0.2 (Pearson Correlation). **(G)** Numbers of highly correlated peaks plotted for each gene (2,284/9,520 expressed genes have at least 1 highly correlated peak). For a given gene, ‘highly correlated’ peaks are those whose R^2^ with gene expression exceeds the 95th percentile among all peaks on the same chromosome. **(H)** Relationships between RNA-seq measurements (top frames, dots represent triplicate RNA-seq measurements/clone) and peak heights (bottom frames, line graphs show individual CREs, colored separately) for select genes.

Founder cells from each clone were profiled by flow cytometry to determine their differentiation state: stem cell memory, effector memory or intermediate (**table S8**). These founder differentiation states manifested in peak variability among their clonal progeny, especially when looking in the first two principal components of peaks (**Fig. S6C**). CRE accessibility around genes linked to differentiation states in vivo also separated the progeny according to founder phenotype indicating a stability of epigenetic features linked to memory T cell differentiation states (**Fig. S6, E and F**).

Interclonal peak differences extended beyond differentiation state. This was particularly visible in two sister clones which were identified in this dataset. These two clones (5a and 5b) descended from memory cells of distinct differentiation states in vivo, yet they generated progeny in vitro with nearly identical protein expression profiles for all markers included in our flow cytometry analysis (**Fig. S6, A and B**). Performing PCA on ATAC-seq peaks, the sister clones separated in PC1/2, reflecting distinct founder states (**Fig. 4B, Fig. S6, C and D)**. In later principal components, however, the two sisters appeared significantly more similar than unrelated clones, again suggesting a heritable layer of clonal identity beneath differentiation state (**Fig. S6D**). Unique patterns of CRE accessibility could be clearly identified for all clones, consistent with highly complex phenotypes defining each clonally expanded population (**Fig. S6G**).

### Chromatin accessibility mirrors interclonal transcriptional differences

We next addressed the extent to which we could ascribe differences in gene expression patterns observed across all clones with variability in chromatin accessibility in our ATAC dataset. A general comparison of ATAC and RNA-seq datasets, revealed that 62% of CREs with interclonal differences were located within 50kbp of genes expressed in our companion RNA-seq dataset (**Fig. 4D**). These variable CREs were evenly distributed between promoters and putative enhancer regions. Looking only at RNA-seq data, we identified 1,899 genes which showed interclonal differences in expression levels (table S9). We identified at least one nearby variable CRE for 1,156 of these genes (RPV>1, 60.8%) (**Fig. 4E**). By correlating CRE and transcriptional differences across a set of 16 clones with high quality data for each measurement we were able to identify 2,934 CRE-gene pairs with correlated activity (Pearson R^2^>0.2) (**Fig. 4F**). This highlights the power of combining both analyses to identify CRE-gene interactions which may be too subtle to identify using a single measurement. As expected, enhancer variability was found to have stronger correlations than promoter variability alone in most cases, although we did find clear evidence of genes with ON/OFF behaviors driven solely by promoter accessibility (**Fig. S7, A and B**). We identified many genes (e.g. *CADM1*, *KLRD1*) with multiple variable CREs linked to transcriptional activity, but most CREs were highly correlated with each other (**Fig. 4G, Fig. S7, C-E**). We found clear evidence for CRE variability linked to ON/OFF patterns of gene expression (e.g. *IL17RB*, *HPGD*, *CADM1*) as well as with tuning transcriptional activity of genes expressed by all cells (e.g. *GNLY*), including many observed in our other datasets (**Fig. 4F, Fig. S7E**). These findings support that clones exhibit diverse, heritable patterns of gene expression and allude to the potential for epigenetic factors to encode a vast array of gene expression profiles among cells of a given type.

### Clonally heritable gene expression in the mouse central nervous system

We next addressed whether our findings extend beyond human lymphocytes. For this we took advantage of TREX, a genetic barcoding approach developed in our lab that enables simultaneous clonal tracking and gene expression profiling in the mouse brain via single-cell transcriptomics (*31*). We delivered a lentiviral barcode library (**Fig. 5A**) into the developing mouse brain at embryonic day 9.5 and isolated barcoded cells from the somatosensory cortex from two mouse brains 14 days post-partum for high-sensitivity, single-cell RNA-seq (Smart-seq3) (**Fig. 5B, Fig. S9**). We obtained a total of 4,010 single cell transcriptome profiles and identified 18 distinct cell types including 7 neuronal, 3 astrocyte, 6 oligodendrocyte and 2 immune cell types (**Fig. 5C, Fig. S10**). We reconstructed 318 multi-cell clones that contained a total of 2,202 cells with an average size of 7 cells per clone (**Fig. S10**).

**Figure 5.**
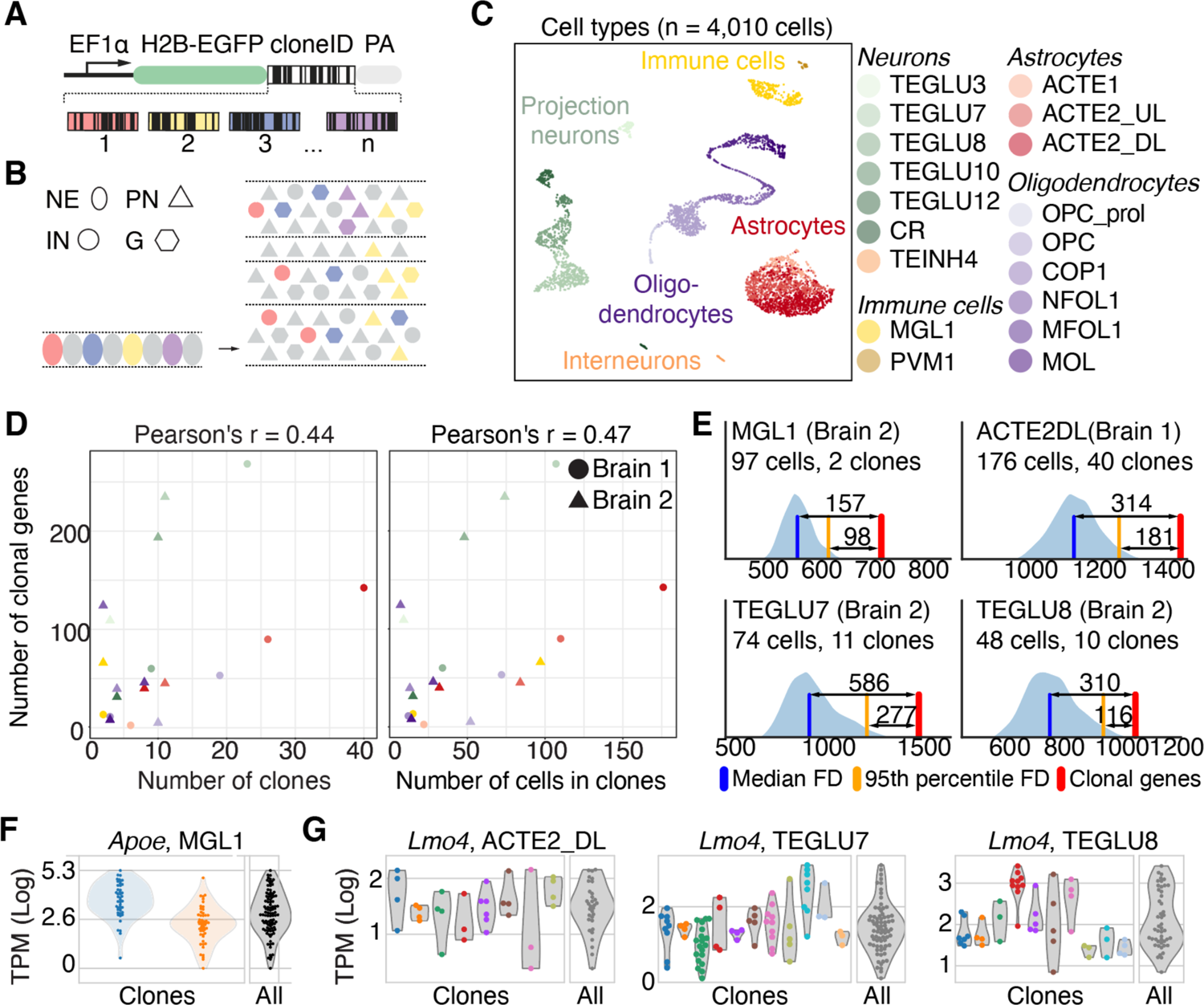
Clonally heritable gene expression in the mouse central nervous system. **(A)** A lentivirus library encoding nuclear-localized EGFP and about 1 million expressed barcodes (‘cloneID’) for unique labeling of progenitor cells and high-throughput clonal tracing. This approach enables simultaneous clonal tracking and gene expression profiling. (**B)** Mouse cortical development from embryonic age 9.5 (E9.5) to post-natal day 14 (P14). Neuroepithelial cells (NE) generate a large diversity of cell types including excitatory projection neurons (PN), inhibitory interneurons (IN) and non-neuronal glia cells (G). Each color represents a distinct barcode. **(C)** Visualization of identified cell classes using UMAP. In total 4,010 single cell transcriptomes were collected from the somatosensory cortex of two P14 mouse brains that were classified into 18 cell types. Capital black letters indicate a unique identifier for each cell type take from www.mousebrain.org. Colors indicate five broader cell type classes: astrocytes (reds), immune (yellows), interneurons (oranges), projection neurons (greens), and oligodendrocytes (purples). **(D)** Scatter plots showing that the number of clonal genes positively correlates with the number of clones (left) and the number of cells in clones (right). **(E)** KDE-smoothed histograms displaying the results of clonal shuffling experiments to identify clonal genes per cell type and brain. Blue KDE-smoothed histogram displays the number of false discoveries (genes with ANOVA F, p<0.05) among 1000 shuffles of clone labels. Blue line is the median number of false discoveries, yellow line the 95th percentile in the count of false discoveries. Red line the number of clonal genes that were found with real clone labels. **(F,G)** Examples of clonally-variable genes *Apoe* **(F)** and *Lmo4* **(G)** in different cell types.

Focusing on populations of cells of the same type where multiple large clones could be identified, we assessed whether we could identify interclonally differentially expressed genes. We found genes with interclonal differences in all cell types and observed that the expression of 12 to 584 genes were differentially expressed between clones of identical cell types (**Fig. S11)**. The number of interclonally differentially expressed genes detected was positively correlated with the number of clones per cell type (Pearson’s r = 0.44) and with the number of cells in clones (Pearson’s r = 0.47) (**Fig. 5D**). The number of genes detected also varied between cell types, e.g. we found 584 clonal genes in layer 2/3 excitatory cortical neurons (TEGLU7, brain 2, 74 cells, 11 clones) and 56 clonally-variable genes in deep layer astrocytes (ACTE2_DL, brain 2, 32 cells, 8 clones) despite similar numbers of clones and number of cells in clones (**Fig. 5E**). Significantly differentially expressed genes included *Apoe* with a wide distribution of graded expression levels in all cortical microglia (MGL1) that could be decomposed into narrow expression patterns between cells belonging to distinct clones (**Fig. 5F**). A differentially expressed gene which was identified in multiple cell types is *Lmo4* which exhibited a range of distinct expression patterns among clones from deep layer astrocytes (ACTE2_DL), layer 2/3 excitatory neurons (TEGLU7) and layer 4 excitatory neurons (TEGLU8) (**Fig. 5G**). Taken together, these data demonstrate that clonally heritable gene expression patterns are present in diverse cell types and species.

## Discussion

Single cell transcriptomics has emerged as a powerful tool to find cellular diversity across developmental stages and tissue types. Diversity is seen even within cell types, where it has been attributed to the stochastic nature of transcription and to fluctuations between transitory cell states. We provide evidence here that heritable transcriptional states provide an additional source of diversity within cell types.

Surveying clonally related cells, we uncovered heterogenous and heritable identities which can persist for more than a year in vivo. The number of distinct phenotypes we see, even within a single cell type, seems limited only by the number of clones we observe. These identities are validated by gene expression signatures, which comprise dozens if not hundreds of independently regulated genes. We provide additional evidence that variable ranges of transcriptional activity between clones lead to corresponding variability in protein expression levels. In large datasets with many different clones, these signatures are strong enough to classify single cells with high accuracy, though they may appear as transcriptional noise without knowledge of clonal structure. However, in datasets with only a few highly expanded clones, the clonally heritable phenotypes emerge as the dominant signal (**Fig. S2**).

We demonstrate that these heritable traits are largely defined by epigenetic features. Interclonal differences in chromatin accessibility in promoter and enhancer regions imparts a wide variety of heritable gene expression states. These include rare expression of genes by a few clones as well as tuned expression of frequently expressed genes, defining specific transcriptional setpoints which differ between clones. The clone-associated variation in expression is neither determined by genetic variation or allele-specific regulation (*17*), and instead likely reflect configurations of regulatory factors (e.g. transcription factors) that can maintain cellular states over longer times. Our findings build on a growing body of evidence that heritable epigenetic features impart long lasting memory of a fixed transcriptional state on differentiated cells (*32*). We would stress, however, that our findings indicate that diversity in heritable epigenetic states could generate heterogeneity within cell types extending far beyond what is conventionally understood to occur during cell type diversification.

This form of heritable cellular diversity may have substantial implications for how we understand the evolution of cellular ecosystems in long-lived multicellular organisms. Variable expression of key genes is known to impart selective advantages in malignant cells in short term assays (*33, 34*) as well as during embryonic development (*35*), and it is possible that similar selective pressures may impact the clonal makeup of tissues under homeostatic or perturbed situations in ageing healthy tissues. This may be particularly important if such variability is not related to short-lived ‘cell states’ but rather reflects the clonal composition of a population. There is now evidence for large expansions of clonally related cells in a variety of different tissues in older humans, suggesting clonal competition is a common feature in aging (*36–38*). Understanding the role that heritable phenotypic diversity plays in this process has the potential to usher in new paradigms for how we view the complex cellular events which contribute to tissue homeostasis, ageing and response to stressors throughout a human lifetime.

## Supplemental Figures

**Fig. S1.**
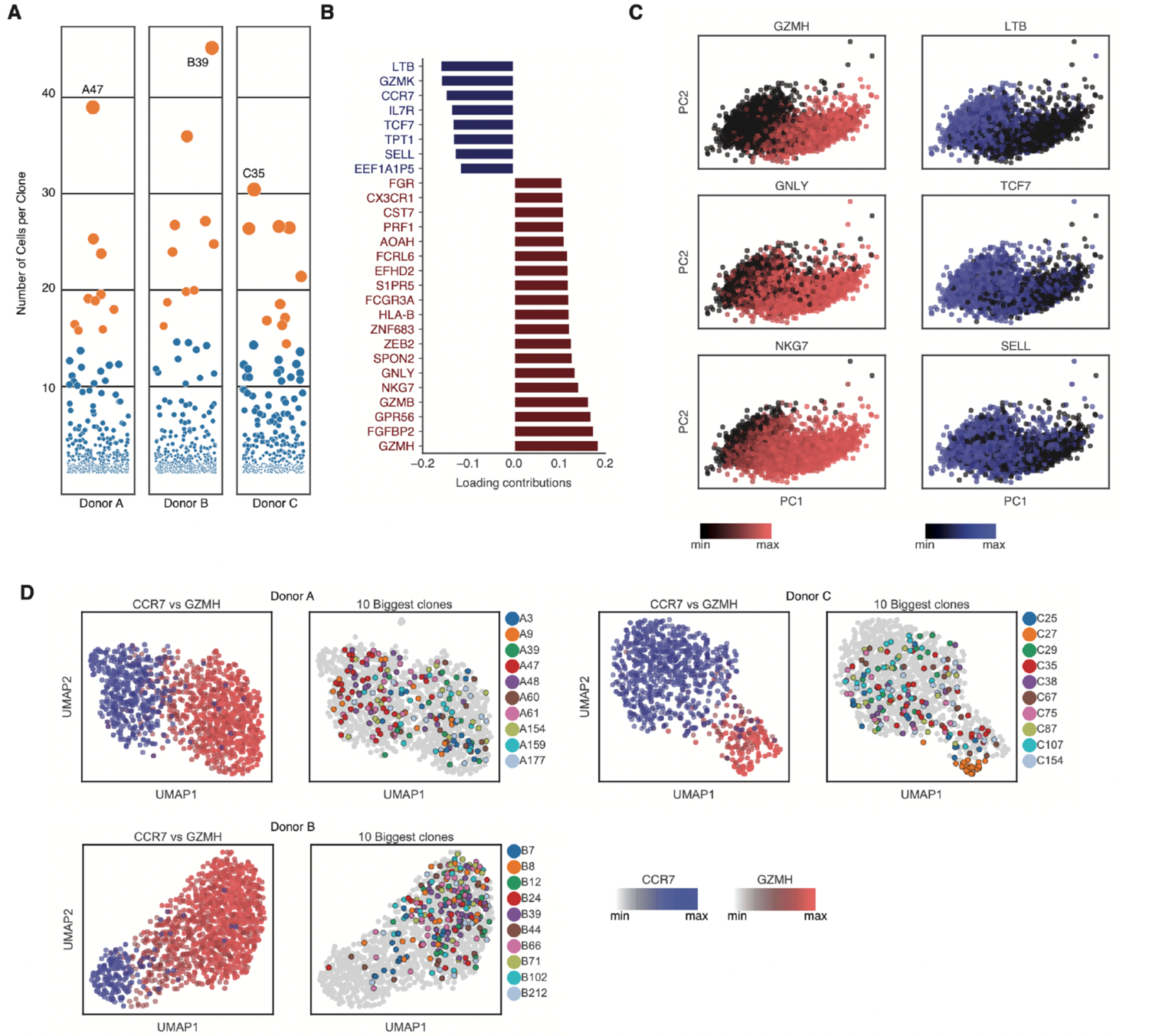
Clonal distribution of memory T cells in vivo relative to differentiation state. **(A)** Clone size distribution for all clones (n> I cell with the same TCRa/b sequences) collected from the YFV antigen-specific circulating memory T cell population of each donor. Dot size and vertical location indicates clone size. Orange dots indicate the top 10 largest clones for each donor. Largest clones are labeled for each donor. **(B)** Top genes contributing to principal component 1, including genes typically associated with highly differentiated cells (red) and less differentiated cells (blue). **(C)** Distribution of all cells (3,837 cells) from all donors based on PC1 and PC2 for all clonal genes. Top genes from PCl are highlighted to show that PC1 orders all cells along a continuum of differentiation states from less differentiated (*LTB, TCF7, SELL* - blue) to more differentiated (*GZMH, GNLY, NKG7* - red) cells. **(D)** UMAPs based on 107 clonal genes showing distribution of all memory cells and top 10 largest clones from each donor. Top genes indicating differentiation states (CCR7-blue, *GZMH*-red) are shown to indicate that UMAP also separates late-stage memory T cells along a continuum based on these genes. Little evidence is seen based on unsupervised UMAP analysis of clonally distinct clustering in UMAP space. Clone 27 in Donor C (orange dots) is an exception as it represents a rare highly differentiated clone in this donor.

**Fig. S2.**
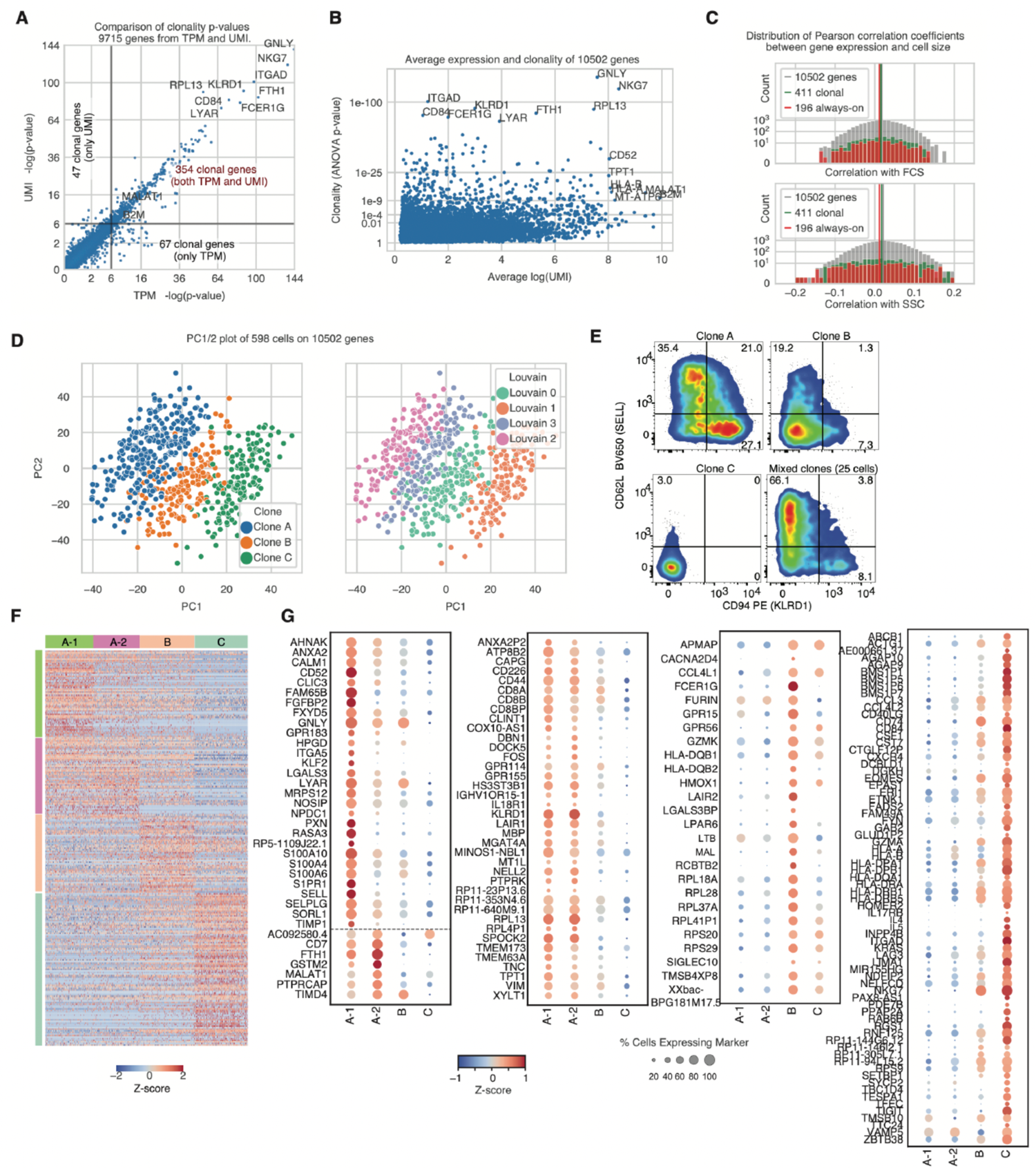
UMI-based assessment of clonal gene expression variability in highly expanded clones. **(A)** Comparison of ANOVA statistics measured for clonal gene expression variability between 3 large clones using either transcripts per million (TPM) or UMI-based quantification strategies for Smart-seq3 data (Table S4). **(B)** Relationship between gene expression levels (based on UMI counts) and clonal variability of gene expression. **(C)** Pearson correlation coefficients for gene expression levels (UMI) vs cell size (FSC, top graph) and cell granularity (SSC, bottom graph) (10,502 detected genes, 411 highly variable clonal genes, or 196 ‘always on’ genes expressed by all cells). **(D)** Principal component analysis (PCA) performed on all genes (10,502 genes) from all high quality single cell libraries (598 cells) from 3 clones. Cells are colored according to clonality.Louvain clustering (right plot) indicating that clusters correspond to unique clones with clone A having two distinct clusters (Louvain 2 and 3, later denoted as A-1 and A-2). **(E)** Protein expression for established differentiation/activation markers on clones A, B, C and a 25-cell mixed clonal bulk population generated in parallel. Clone A shows a clear split in the population according to these two markers which typically are associated with more highly differentiated cells (CD94, gene ID: *KLRDJ)* and less differentiated cells (CD62L, gene ID: *SELL).* **(F)** Heatmap highlighting clonally distinct gene expression profiles, 41 l genes and 598 cells. (G) Among 411 genes, those which identify each clonal population including genes which separate sub-clonal populations A-1 and A-2 (left) and genes enriched in Clone A vs B or C, and genes enriched in Clone B vs A or C, and genes enriched in Clone C vs A or B.

**Fig. S3.**
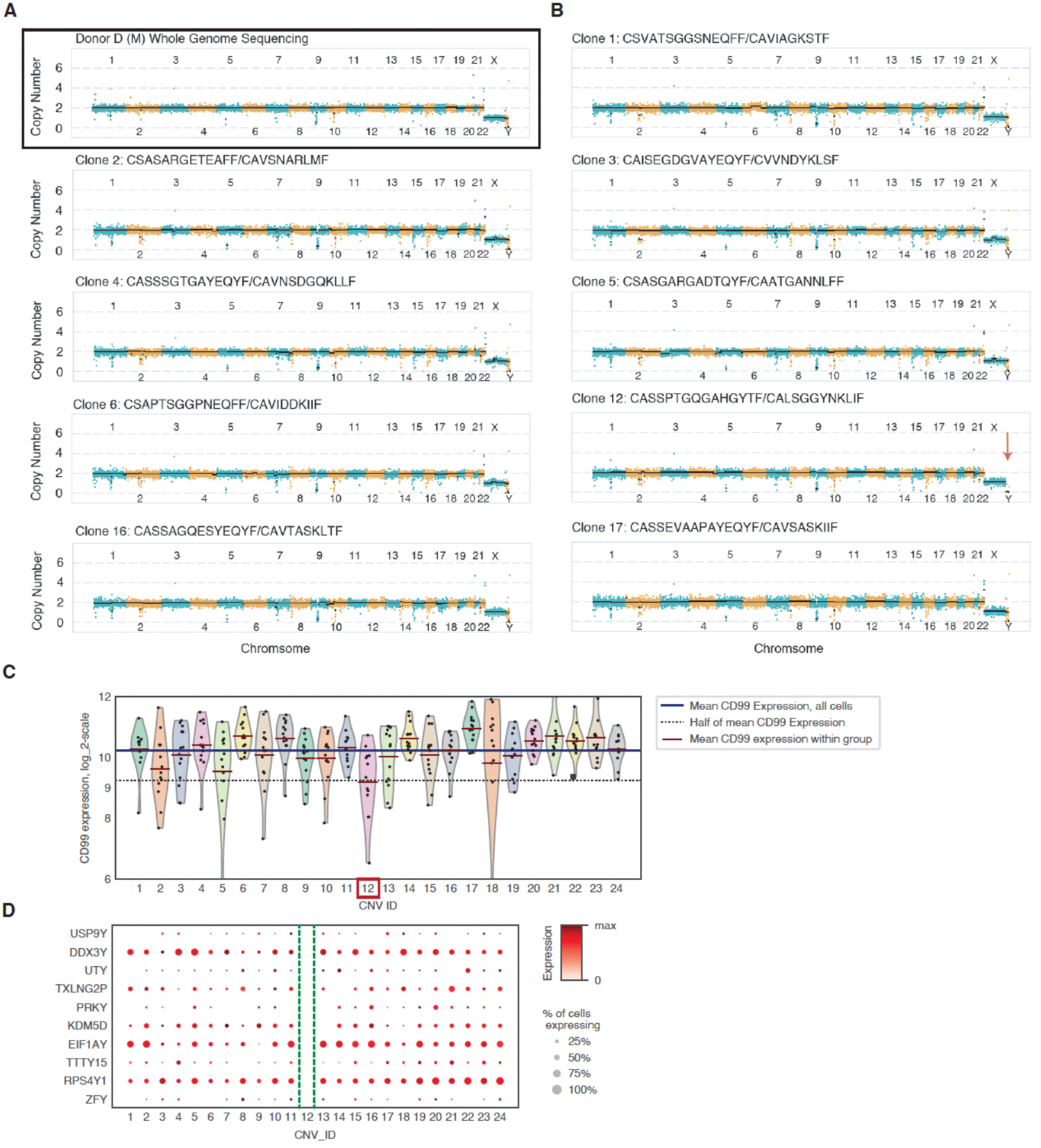
Copy number variations are uncommon in cloually expanded lymphocytes. **(A)** CNV analysis of Donor D based on whole genome sequencing perfmmed on gDNA isolated from pe1ipheral blood (30x coverage downsampled to match single cell datasets). **(B)** Clonal CNV analysis for 9 (out of 24) clonal populations analyzed in figure 3 (sister clones are analyzed separately). The 11 clones not shown displayed no obvious CNVs and were omitted for space reasons, all data is deposited together. The clonal numbers and TCR sequences (TCRBfrCRA) are shown for each displayed population. Clone 12 is highlighted due to a clear loss of Y-chromosome (LOY) obse1ved in our analysis (red a.now). CNVs were quantified based on SOOkbp bins so small CNV are unlikely to be detected in this analysis. **(C)** Matched single cell RNAseq analysis of clones shows clonal vaiiability around CD99 which is expressed on both the X and Y chromosome. Clone 12 has on average a 50% reduction in CD99 expression consistent with a loss of a single copy of this gene. **(D)** Complete loss of expression of remaining Y-chromosome genes for clone 12. CNV_ID matches deposited data and individual sister clones are analyzed as separate datasets in this analysis.

**Fig. S4.**
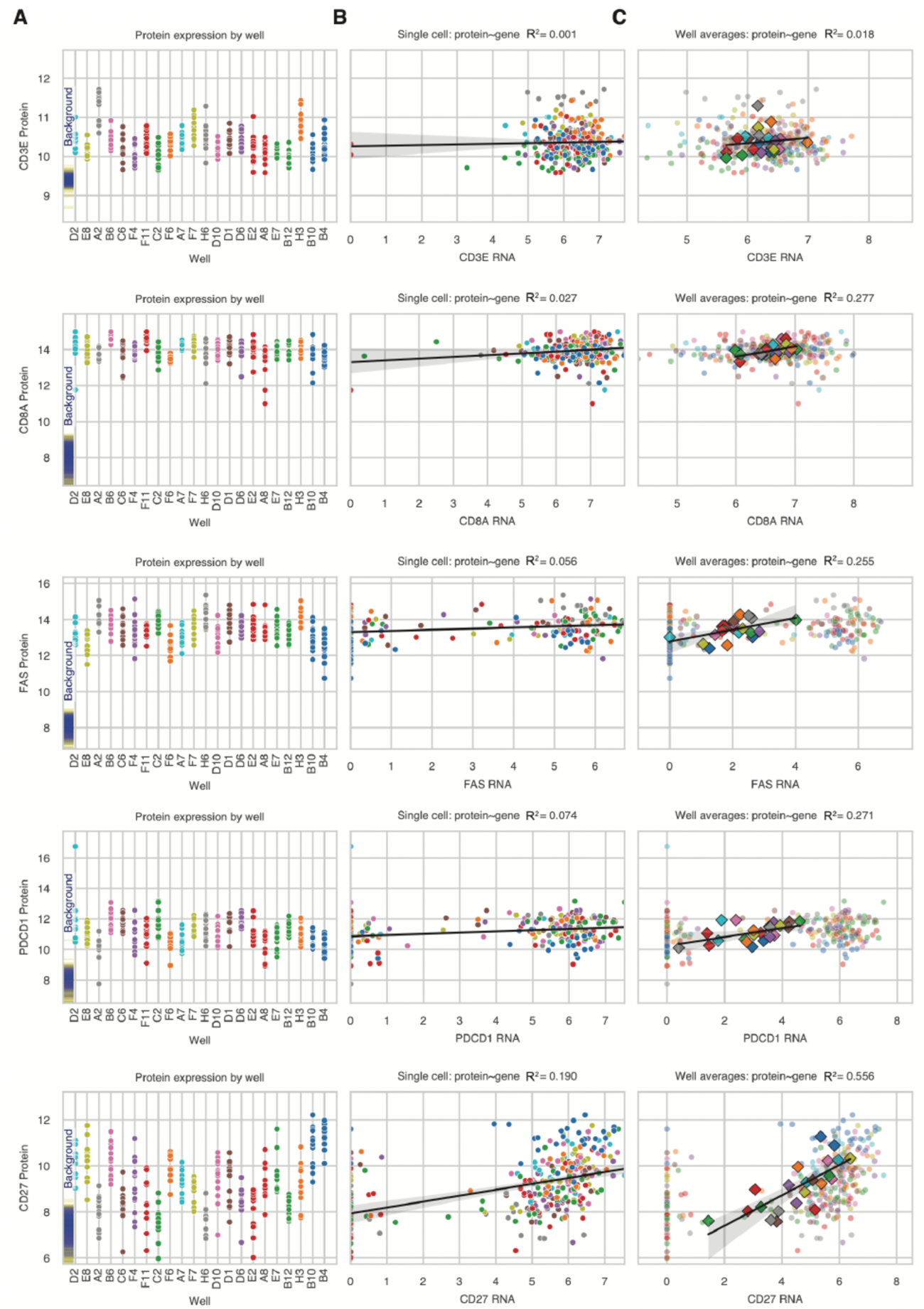
Clonally variable gene expression is mirrored in protein expression levels and correlations between each measurement are strengthened by measuring clonal averages. **(A)** Protein expression levels measured on single cells during index sorting for clones from Fig. 3 (Donor D, Day 536 post-vaccination, Table S6). Clones are labeled by well in which they were expanded and sister clones are plotted separately at the end of the graph (C11/wells A8:E2 - red; C113/wells BI 2/E7 - green, Cl54/wells 810/84 - blue). Well H4 contained single cells belonging to both clones 11 and 54 and was omitted in this analysis. Expression levels oflineage specific proteins (CD3E, CD8A) and differentiation/activation proteins (CD27, FAS, PD-I *(PDCDJ))* are shown separately. Background levels of detection represent autofluorescence intensity for each channel in control cells stained in parallel with all markers except the indicated marker (fluorescence minus one (FMO) controls). **(B)** Correlations of matched single cell RNA-seq gene expression measurements (TPM values) for each cell with protein expression measurements from panel a. Most markers show weak correlations (coefficient of determination R2<0.1) with the exception of CD27 (R2=0.19). **(C)** Average clonal single cell RNA-seq measurements correlated with average clonal protein expression measurements reveals heightened correlations between RNA and protein across clonal populations. We interpret this to reflect a more accurate measure of the range of variance for each variable which can be temporally uncoupled at the single cell level.

**Fig. S5.**
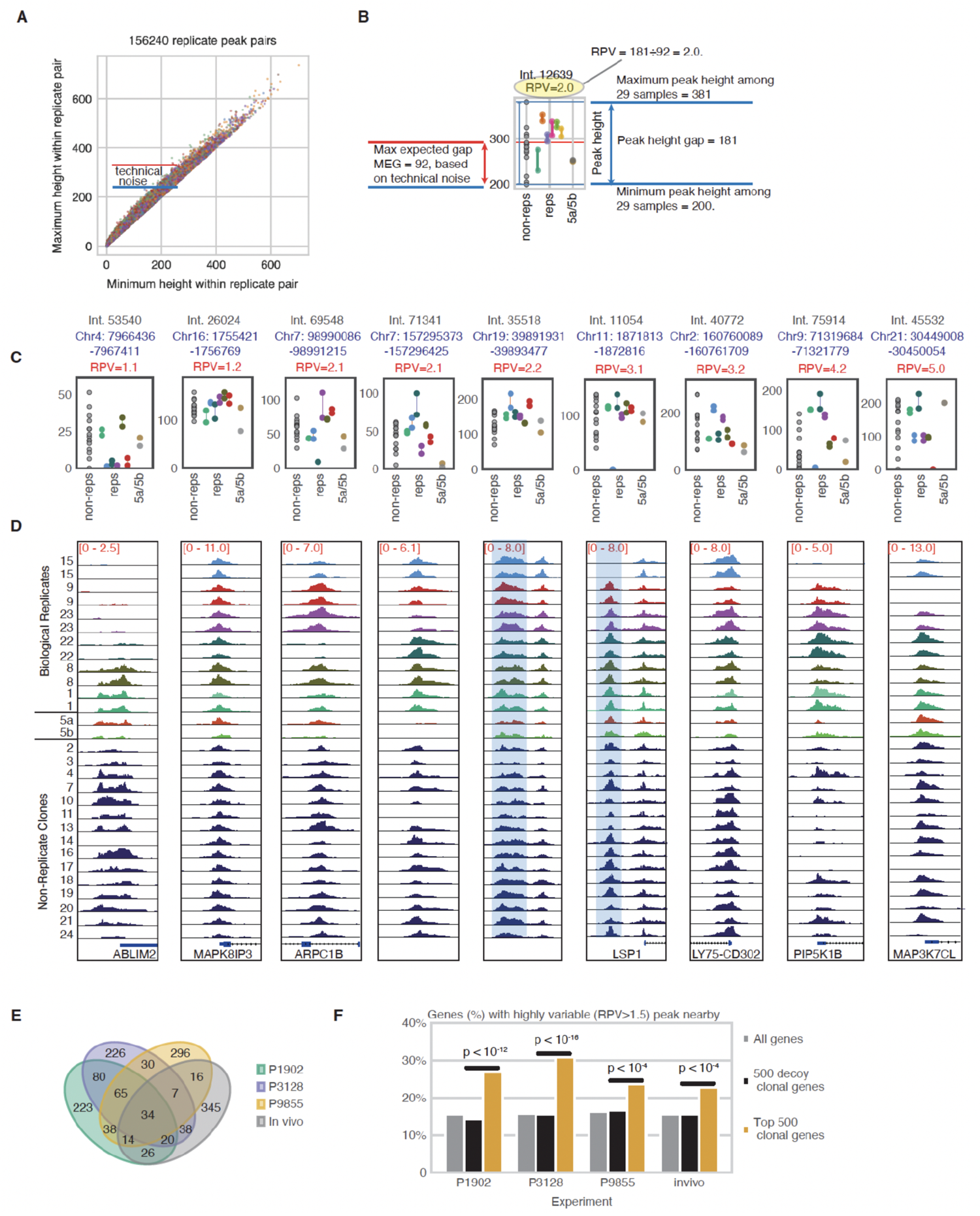
Description of clonal CRE variability metrics and examples of variable peaks according to different peak variability scores. **(A)** Technical ‘noise’ associated with assay measurement s. 26,040 CREs and 12 samples in 6 replicate pairs yield 156,240 replicate peak pairs (RPPs). For each RPP, smallest height potted on x and largest on y. Difference represents technical noise. **(B)** Plot in (A) is used to estimate the maximum expected gap (MEG) one might expect for a given CRE due to noise. Relative peak variability of a CRE is the quotient of actual gap (maximum height among 29 samples minus minimwn height) by the MEG (swnmary of all peaks in Table S7). **(C)** Examples of peak height variability for single clones (gray, non-reps), biological replicates (reps), and sister clones (5a/5b). Examples are ordered according to RPV score and y-axis scales vary between examples. **(D)** Matched plots from Integrated Genome Viewer (IGV, Broad Institute) for each peak in panel (C). (E) Overlaps between the top clonal genes (approx. 500) from each dataset. Approximately half of the clonal genes identified in the ATAC/RNA experiment (Fig 4) are also identified in our other datasets (Fig. 2 = Pl902, Fig. 3 = P3128, Fig. 4 = P9855) (F) The % of genes in each dataset with a nearby peak showing high clonal variability (RPV> 1.5). ‘Decoy genes’ are generated by applying the same statistical test for interclonal variability to shuffled clones. A binomial test was performed to compare the proportions of genes among the interclonally variable genes and among the decoy genes (one sided p-values are reported).

**Fig. S6.**
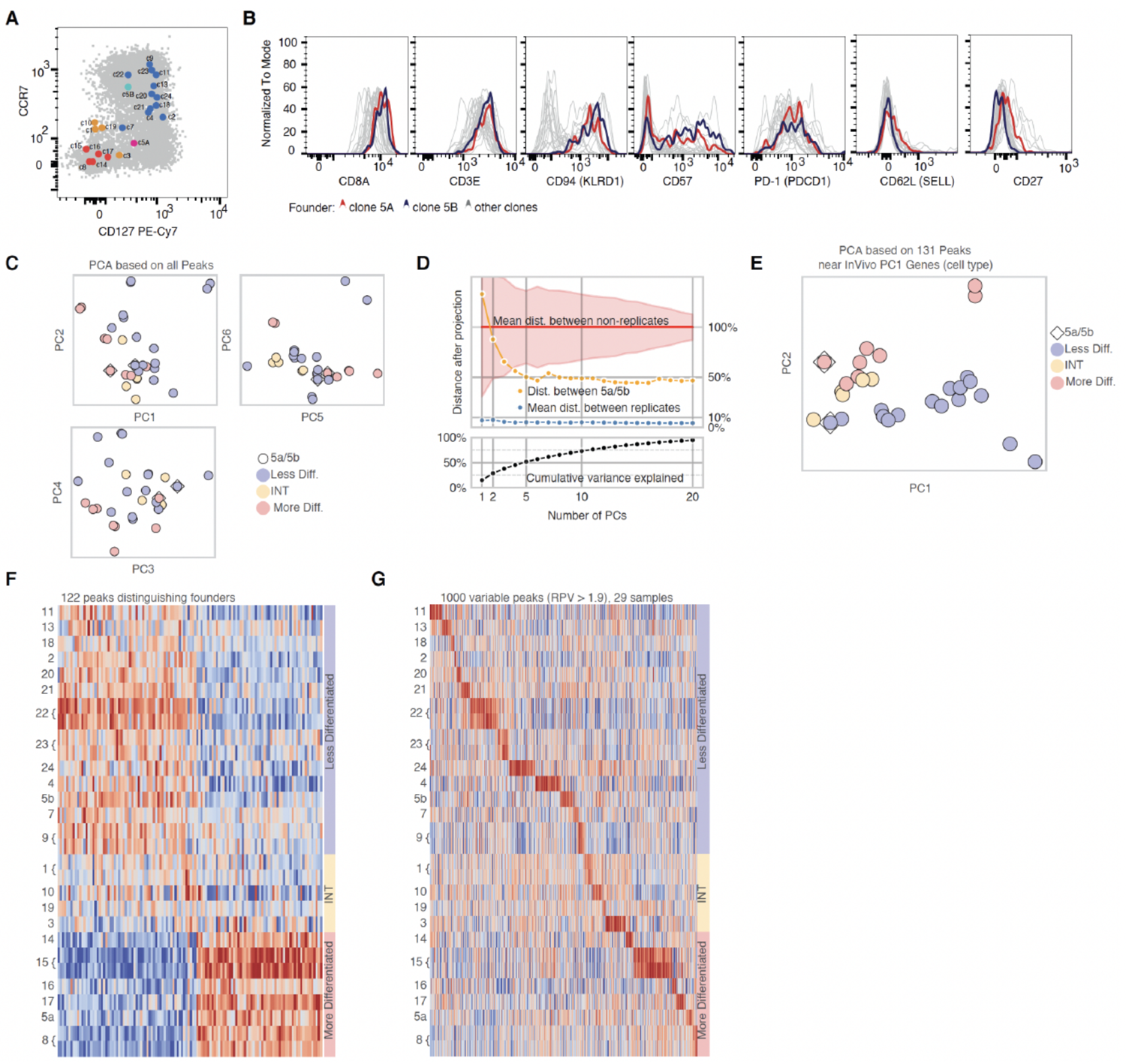
Clonal maintenance of parental chromatin expression states. **(A)** Memory T cells sorted from Donor E to generate datasets in figure 4 are labelled on a scatter plot showing total distribution of all CDS+ T cells. Markers of differentiation state are included to distinguish clones occupying different differentiation states in vivo. Less differentiated clones (blue, top right) express high levels of CCR? and IL7R (both contribute significantly to PCl in Fig. SIB, **C)**. More differentiated clones are negative for these markers and colored either yellow (intermediate) or red (most differentiated). Each population is labelled according to standard nomenclature in the CDS+ T cell field: Less differentiated - SCM (stern cell memory), Intennediate - INT, and more differentiated - EM (effector memory). Sister clones (Sa and Sb) were found to occupy distinct memory differentiation states in vivo (Sa = EM, pink and Sb = SCM, light blue). **(B)** Despite sisters having different founder phenotypes, progeny had remarkably similar protein expression levels for all activation and differentiation markers profiled. **(C)** Clonally expanded progeny displayed in the first six principal components based on all 26,040 peaks. Founder identity contributes to differences in the first few PCs where it separates sister progeny (shown in boxes in each plot). Sisters appear closer in PCS,6. **(D)** Distance between unrelated clones (shaded area, all clones range), replicate clones (blue), and sisters (yellow) according to different PCs ranging from PC1-PC20. Black line indicates contribution of each PC to total variance of peak heights in dataset. **(E)** PCA performed on all clones labelled based on founder identity using peaks nearby genes contributing to memory T cell differentiation differences in vivo (fig SIB, PCl genes). Note Sa and Sb separated to their appropriate founder group. **(F)** Progeny of SCM vs EM memory T cells have clear signatures based on their clonal ancestry with INT cells exhibiting intermediate identities. g, Clones exhibit specific enrichment for sets of CRE independent of founder identity.

**Fig. S7.**
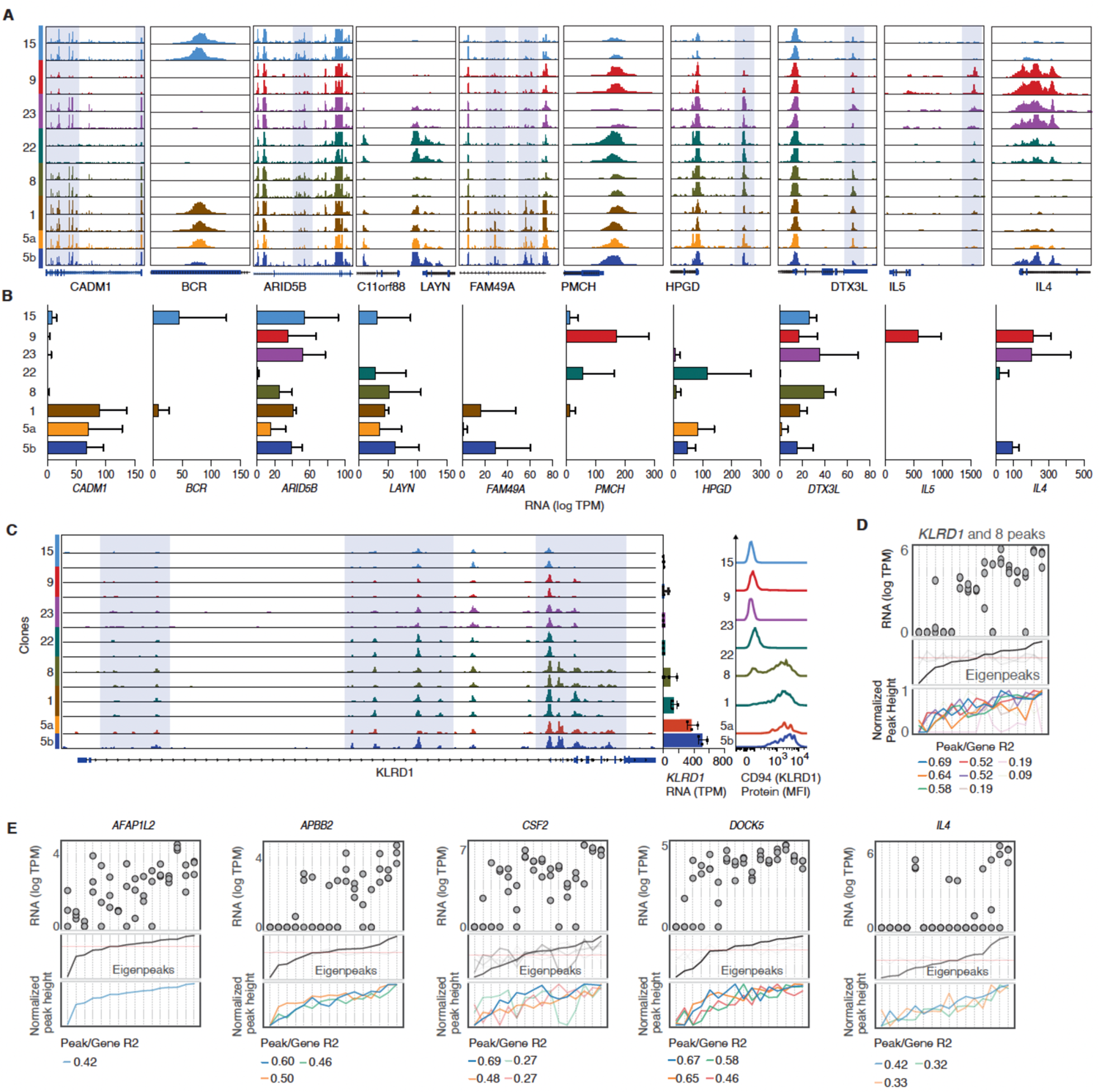
Linking CRE variability to clonal gene expression variability. **(A)** Examples of different CREs with vaiiable accessibility in a variety of genes with different fimctions. Clonal biological replicates and sister replicates are shown (non-replicates not shown for space). Insome cases peaks are highlighted when multiple peaks are shown to indicate peaks most associated with gene expression variability. **(B)** corresponding RNA-seq gene expression measurements for nearby genes to CRE in panel a. Clones are color coded to match panel a. Bars represent the average and standard deviation for niplicate RNAseq measurements (2S cells/replicate) for each gene (TPM values). Sister clones (yellow and blue) each are measured separately while clones with biological replicates for ATAC-seq only have 3 RNAseq measurements/clone. **(C)** Example of a gene *(KLRDJ)* with a Iarge munber of conelated peaks (S peaks with R2>0.S) showing entire gene u·ack from the TSS to TIS. Clonal biological replicates and sister clones are shown as in panel a. High­ lighted regions show clonally vaiiable peaks. Panels on the light show mRNA (TPM) expression levels and protein (MFI) expression levels for each clonal population demonsu·ating increasing levels relative to peak height va1iability. Note that all cells for clone I are CD94 *(KLRDI* protein) positive yet mRNA levels are neai-Jy 2-fold lower and CRE accessibility is reduced relative to clones Sa and Sb. **(D)** A summa1y plot showing highly COITelated peaks and mRNA expression for *KLRDI* across 16 clonal populations showing gradual increases in expression nmed by CRE accessibility. We inn·oduce ‘eigenpeaks’ (Methods) as a ctunulative effect of all CRE activity to describe *KLRDJ* expression. Eigenpeaks are calculated solely from CRE covariance (eigenvectors of peak covariance matlix) and independently of mRNA expression. Clones are ordered based on eigenpeak va lues revealing cleat·relationships to mRNA expression (top panel). **(E)** Examples of genes ordered by eigenpeak values demonstrating that coordinated activities of many CREs can tune gene expression levels in clonally distinct patterns.

**Fig. S8.**
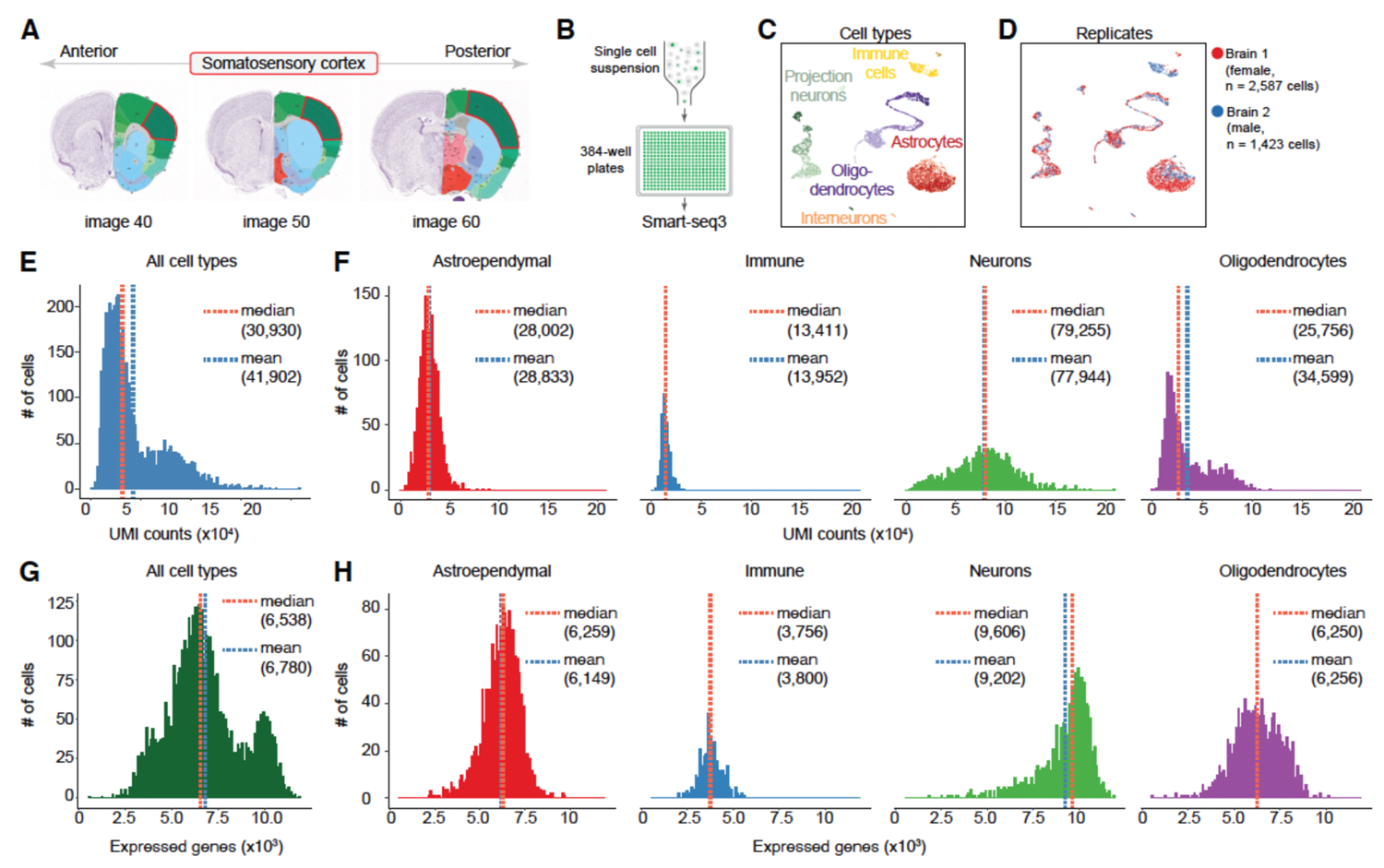
Dissection strategy, replicates and expression metrics for the mouse brain dataset. **(A)** Each brain was cut using a 1 IIl1Il coronal brain slicer and brain regions shown in red contour along the ante1ior/poste1ior axis were dissected for tissue dissociation. Image m1mbers refer to the image m1mber from the Allen Brain Atlas (http://atlas.brain-map.org/atlas?atlas=l#atlas=l). **(B)** Sections from two individual brains were dissociated separately and single EGFP+ cells were sorted into 384-well plates followed by library prep using Smart-seq3. c, d, UMAP visualizations grouped by major cell types **(C)** or replicate **(D)**. **(E, F)** Number of transcripts (unique molecular identifiers, UMis) per cell for the entire dataset (E) and for each major cell type (F). (G, H) Number of expressed genes per cell for the entire dataset (G) and for each major cell type (H).

**Fig S9.**
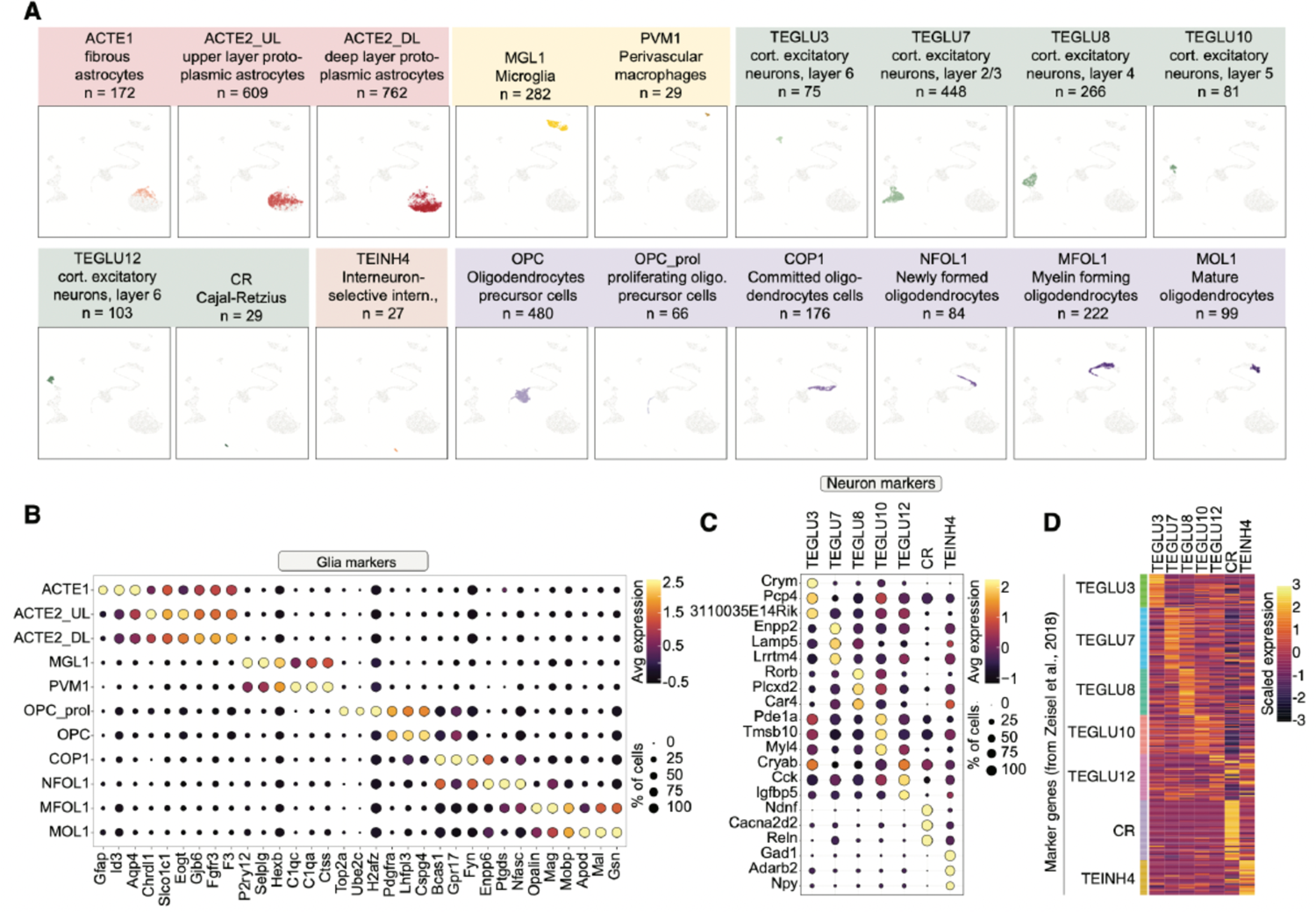
Cell type classification and markers in mouse brain datasets. **(A)** Separate UMAP visualizations for all cell types and corresponding number of cells per type identified in this study. Colors indicate five major cell type classes: astroependymal (reds), immune (yellows), interneurons (oranges), projection neurons (greens) and oligodendrocytes (purples). We followed the nomencla­ ture from Zeise! et al., 2018 to annotate cell types and further subdivided ACTE2 into deep layer (DL) and upper layer (UL) cells as described by Bayraktar et al., 2020. **(B,** _C)_ Gene expression of markers for each glia _(C)_ and for each neuronal cell type **(D).** For each cell type the top three marker genes were identified, and unique genes plotted as dot plots. Expression values represent scaled average gene expression per cell type. **_(D)_** Heatmap showing the expression for unique differentially expressed genes (rows) identi­ fied in a publi shed mouse brain atlas in each of the corresponding clusters (columns) identified in this study.

**Fig. S10.**
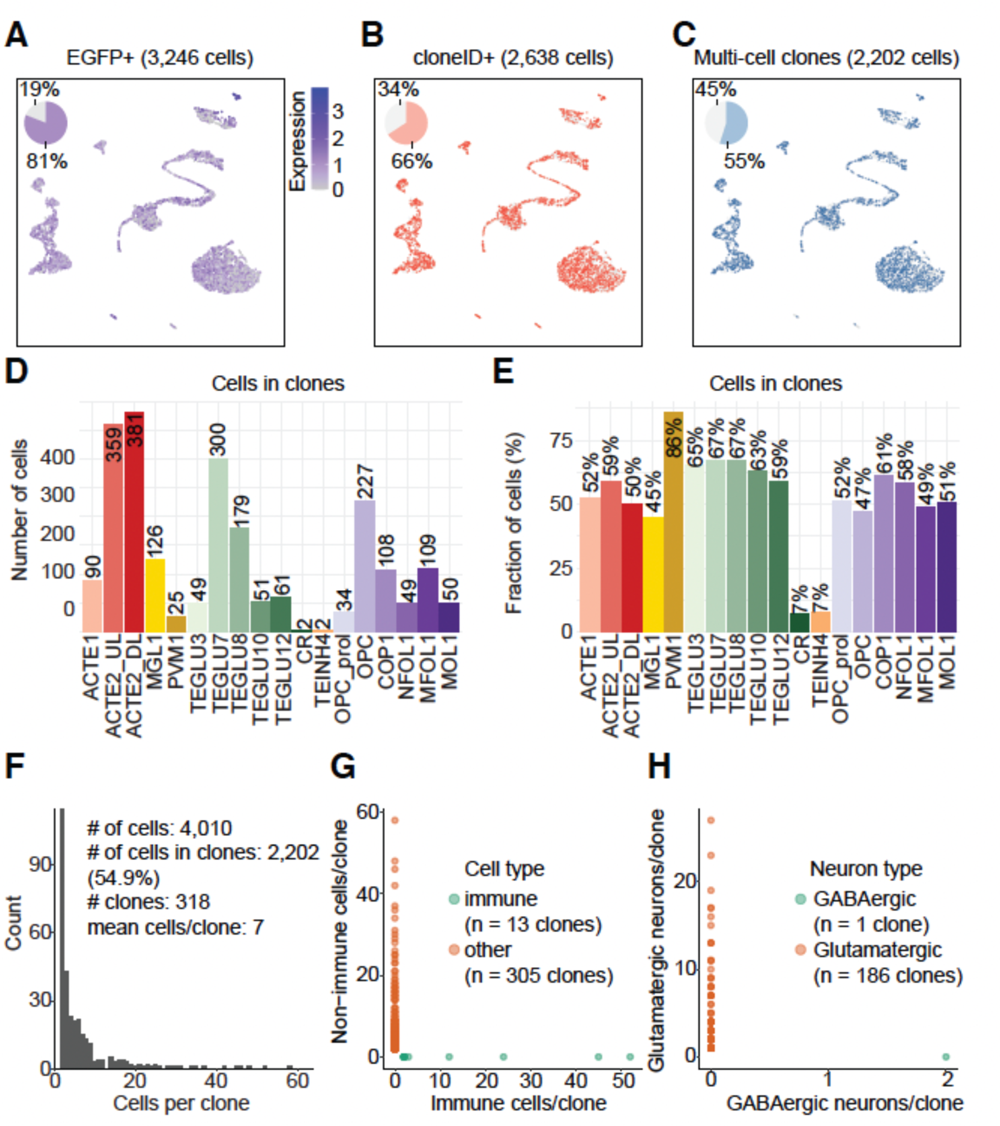
Clone reconstruction and clonal composition for the mouse brain dataset. **(A)** UMAP embeddings and normalized EGFP expression levels in all cells (n = 4,010) isolated from postnatal mouse brains that were injected with EFla-H2B-EGFP-cloneID libraries at E9.5 (grey dots). A total of 3,246 cells contained at least one EGFP transcript (blue). **(B, C)** A total of 2,638 cells contained a cloneID (B, red) and 2,202 cells were contained in multi-cell clones (C, light blue) defined as groups of minimum two cells that share the same cloneID. **(D)** Total number of cells in multi-cell clones for each cell type. (E) Fraction of cells per cell type found in multi-cell clones. **(F)** Histogram and key summaiy metrics showing the clone size distribution for all reconstructed clones. **(G)** Scatter plots showing the number of cells in clones containing immune cells (x-axis, green) or non-immune cells (y-axis, red). **(H)** Scatter plots showing the number of cells in clones containing GABAergic inhibito1y neurons (x-axis, green) or glutamatergic, excitato1y cells (y-axis, red). We never observed a shared cloneID between cell types de1ived from different progenitor cells (immtme and non-immune cells, inhibitory and excitat01y neurons) indicating that our clone calling pipeline is enor-free.

**Fig. S11.**
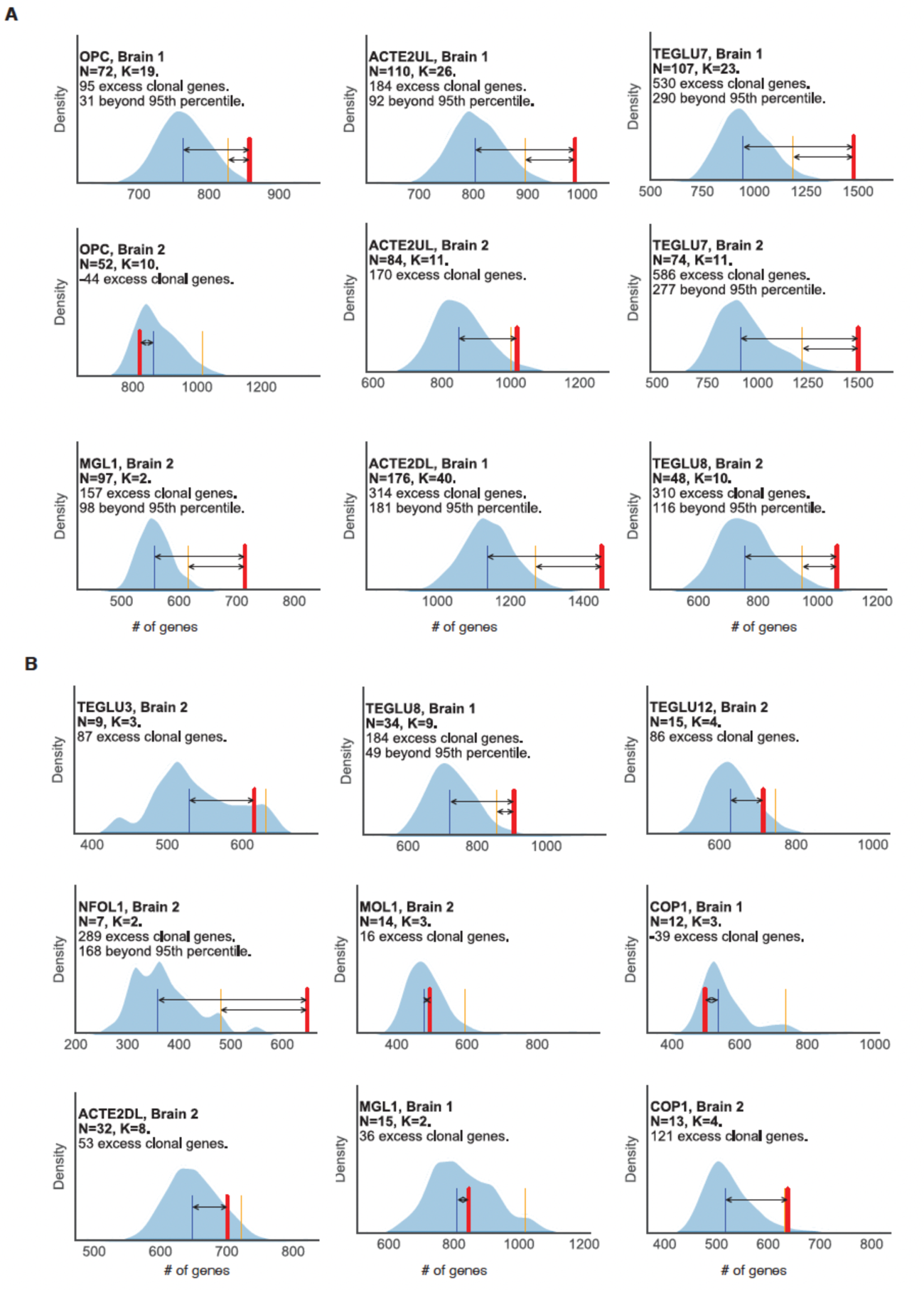
Clonally variable genes incell types of the mouse central nervous system grouped by clone size/number (A, B) KDE-smoothed histograms displaying the results of clonal shuffling experiments to identify clonal genes per cell type and brain (see Methods for details). **(A)** Cell types where clones were found with more than 40 cells (in total) and **(B)** cell types with clones having less than 40 cells in total. N= the number of different clones detected for each cell type and K= the number of total cells found in all clones. The blue line is the average false deteection rate and the orange line is the *_9th_* percentile for false detection with shuffled clones. The red line indicates the number of significant genes detected in true clonal populations.

## Acknowledgements

We wish to thank members of the Frisén and Michaelsson laboratories for assistance in sample collection and preparation of the final manuscript. In particular we would like to thank Sarantis Giatrellis for help with flow cytometry and the sequencing core facility at SciLife Labs for help with sequencing and data management. We thank the following funding agencies for providing support for this study: This study was supported by grants from the Swedish Research Council, the Swedish Cancer Society, the Karolinska Institute, the Strategic Research Programme in Stem Cells and Regenerative Medicine at Karolinska Institutet (StratRegen), the Swedish Society for Strategic Research, and Knut och Alice Wallenbergs Stiftelse.

## Author Contributions

J.E.M., J.M., M.W., and J.F. conceived and designed the study. J.E.M. and J.M. isolated all human cells and performed all related experiments. M.R. and M.H.J. performed all mouse experiments. J.E.M., M.W., M.R., M.H.J., J.H., C.J.E., L.B., M.M. performed data analysis with the supervision of R.S. J.M. and J.F.. H.T. and J.B. processed sequencing data with the supervision of J.La. and J.Lu.. J.E.M., M.W., J.M. and J.F. wrote the manuscript with assistance from all other authors. M.W. generated all computer code and maintains the GitHub repository containing information for processing data.

## Materials and Methods

### Data Generation

#### Human study subjects

HLA-A2+ human volunteers were identified from an ongoing study examining the longitudinal immune response to yellow fever vaccine YFV-17D (approved by the Regional Ethical Review Board in Stockholm, Sweden: 2008/1881-31/4, 2013/216-32, and 2014/1890-32). Written informed consent was given by all participants prior to study start. Longitudinal venous blood samples were collected in BD vacutainer tubes with heparin (BD Biosciences) and total peripheral blood mononuclear cells (PBMCs) were isolated by density centrifugation according to the manufacturers protocol (Lymphoprep, Stem Cell Technologies). Samples were cryopreserved at a concentration of 10^7^ cells per milliliter in a solution of 90% fetal bovine serum (FBS, Gibco) and 10% dimethylsulfoxide (SigmaAldrich) and stored in liquid nitrogen for later use.

#### Isolation of antigen specific CD8+ T cells from total PBMCs

Cryopreserved samples were rapidly thawed at 37°C and washed in FACS buffer (PBS supplemented with 2% FBS and 2mM EDTA). CD8+ T cells were isolated by negative selection using magnetic beads following the instructions from the manufacturer (Miltenyi Biotec). The purified CD8+ T cells were incubated with APC-conjugated HLA-A2/YFV NS4b (LLWNGPMAV) dextramer (Immudex) for 15 min at 4°C, followed by addition of anti-CD3e Alexa700 (clone UCHT-1, BD Biosciences), anti-CD8a APC-Cy7 (clone SK1, BD Bioscience), anti-CD14 V500 (clone φP9, BD Biosciences), anti-CD19 V500 (clone HIB19, BD Biosciences), and Live/Dead Aqua dead cell stain (ThermoFisher) for an additional 15min at 4°C. Cells were washed twice in FACS buffer and suspended in FACS buffer for sorting.

Single, HLA-A2/YFV NS4b-dextramer+, live, CD14-CD19-CD3+CD8+ cells were sorted on single cell sort mode with index sorting. For the experiment where ATAC and RNA-seq were performed on the progeny clones, we used a more comprehensive antibody panel for sorting: anti-CD3e PE-Cy5 (clone UCHT1, BD Biosciences), anti-CD8a Alexa700 (clone SK1, Biolegend), anti-CD14 V500 (clone φP9), anti-CD19 V500 (clone HIB19), anti-CCR7 BV421 (clone G043H7, BioLegend), anti-CD45RA PE-Cy5.5 (clone MEM-56, ThermoFisher), anti-CD127 (clone A019D5, BioLegend), anti-CD62L (clone DREG-56, BioLegend), anti-CD57 FITC (clone NK1, BD Biosciences), anti-KLRG1 APC-Fire750 (clone SA231A2, BioLegend), anti-CD26 PE-CF594 (clone M-A261, BD Biosciences), anti-CD94 PE (clone DX22, BioLegend), Live/Dead Aqua dead cell stain (Invitrogen). Founder cells were classified according to expression of CCR7 and CD127, where stem cell memory (SCM) T cells were defined as CCR7^high^CD127^+^ and effector memory (EM) T cells as CCR7^low/-^CD127^-^. Intermediates (INT) between SCM and EM were CCR7^low^ and/or CD127^+^. Virtually all founder cells expressed CD45RA, and only EM expressed CD57. A summary of normalized protein expression (log2) for each clone can be found in Table S9.

#### Clonal Expansion of YFV-specific CD8+ T cells in vitro

Single live CD8^+^CD3^+^HLA-A2/YFV-dextramer^+^ cells were index sorted directly into 96 well U-bottom plates containing 2 μg/ml YFV NS5b peptide (LLWNGPMAV, JPT Peptide Technologies), 20U/ml IL-2, and 50.000 irradiated (40Gy) CD3-depleted autologous PBMCs in T cell media (RPMI1640 with 10% heat inactivated human AB sera, 1mM <u>sodium pyruvate</u>, 10mM <u>HEPES</u>, 50 μM 2-mercaptoethanol, 1mM L-glutamine, 100U/ml penicillin and 50 μg/ml streptomycin) and were cultured for 18-22 days. Every 4–5 days half of the media was replaced with fresh T cell media containing 50U/ml IL-2 and 2 μg/ml peptide. After 18-21 days, 10% of the cells of the cells from each well were mixed with 10μl AccuCount particles (Spherotec), stained with anti-CD3e Alexa700, anti-CD8a APC-Cy7, anti-CD14 V500, anti-CD19 V500 (BD Biosciences), and Live/Dead Aqua dead cell stain (ThermoFisher), and analyzed on a BD Fortessa flow cytometer (BD Bioscience) to detect and enumerate expanded live CD3+CD8+ T cells. After identifying all wells containing expanded HLA-A2/YFV NS4b-specific CD8+ T cell clones, individual clones were selected for downstream analysis based on having larger expansions. Selected clones were washed in FACS buffer and stained with the same panel as described above (“*Isolation of antigen specific CD8+ T cells from total PBMCs*”), and in addition with anti-CD27 BV786 (clone L128, BD Bioscience), anti-PD1 BV711 (clone EH12.1, BD Biosciences), anti-CD94 PE (clone DX22, BioLegend), anti-CD62L BV650 (clone DREG-56, BioLegend), and anti-CD57 FITC (clone NK1, BD Biosciences).

#### Lentivirus barcode libraries

For mouse experiments, lentivirus preparations have been used as described previously (*31*). Briefly, plasmid libraries encoding a 30N random barcode (“cloneID”) downstream of an H2B-EGFP transgene driven by the human EF1a promoter (EF1a-H2B-EGFP-30N) were generated using Gibson assembly. Plasmid cloneID libraries were used for virus particle production by GEG-Tech (Paris, France) and viruses with a titre of 1.27 x 10^9^ TU/ml were used for all applications. A typical lentivirus preparation contained about 1.57×10^6^ cloneIDs/µl with a largely uniform representation and high sequence diversity (*31*).

#### Mice

CD-1 mice obtained from Charles River Germany were used for all experiments. Animals were housed in standard housing conditions with 12:12-hour light:dark cycles with food and water ad libitum. All experimental procedures were approved by the Stockholms Norra Djurförsöksetiska Nämnd.

#### Ultrasound-guided in utero microinjection

To target the developing mouse nervous system, timed pregnancies were set up overnight, plug positive females were identified the next morning and counted as embryonic (E) age 0.5. Ultrasound check was performed at E8.5 to verify the pregnancy. Pregnant females at E9.5 of gestation were anaesthetized with isoflurane, uterine horns were exposed, each embryonic forebrain injected with 0.6µl of lentivirus corresponding to 0.94×10^6^ unique cloneIDs (*31*) and 4-8 embryos injected per litter. Surgical procedures were limited to 30 min to maximize survival rates.

#### Single-cell dissociations of brain tissue and flow cytometry

Two mice with an age of 2 weeks (postnatal day 14, P14) were sacrificed with an overdose of isoflurane, followed by transcardial perfusion with ice cold artificial cerebrospinal fluid (aCSF: 87 mM NaCl, 2.5 mM KCl, 1.25 mM NaH_2_PO_4_, 26 mM NaHCO_3_, 75 mM sucrose, 20 mM glucose, 2 mM CaCl_2_, 2 mM MgSO_4_). Mice were decapitated, the brain was collected in ice-cold aCSF, 1 mm coronal slices collected using an acrylic brain matrix for mouse (World Precision Instruments) and the primary somatosensory cortex from three slices per brain (Fig. S8) was microdissected under a stereo microscope with a cooled platform. Tissue pieces were dissociated using the Papain dissociation system (Worthington Biochemical) with an enzymatic digestion step of 20-30min followed by manual trituration using fire polished Pasteur pipettes. Dissociated tissue pieces were filtered through a sterile 30 µm aCSF-equilibrated Filcon strainer (BD Biosciences) into a 15 ml centrifuge tube containing 9 ml of aCSF and 0.5% BSA. The suspension was mixed well, cells were pelleted in a cooled centrifuge at 300g for 5 min, supernatant carefully removed, and cells resuspended in 1 ml aCSF containing reconstituted ovomucoid protease inhibitor with bovine serum albumin. A discontinuous density gradient was prepared by carefully overlaying 2 ml undiluted albumin-inhibitor solution with 1 ml of cell suspension followed by centrifugation at 100g for 6 minutes at 4°C. The supernatant was carefully removed, the cell pellet resuspended in 1 ml aCSF containing 0.5% BSA and the cell suspension transferred to a round bottom tube (BD Biosciences) for flow cytometry. Single EGFP+ cells were sorted on a BD Influx equipped with a 140 µm nozzle and a cooling unit with a sample temperature of 4°C and collected into 384-well plates (Armadillo) for Smart-seq3 as described above.

#### Single Cell RNA sequencing of T cells and mouse brain cells

Both Smart-seq2 and Smart-seq3 protocols were used to generate single cell RNA-seq libraries from both *in vivo* and *in vitro* expanded HLA-A2/YFV NS4b-specific CD8+ T cells. Smart-seq3 was used to generate libraries for mouse central nervous system cells.

For ***Smart-seq2*** labeled T cells were sorted into 96-well V-bottom plates (Thermo) containing lysis buffer (0.1%Triton X-100, 2.5mM dNTP, 2.5μM Oligo-dT, 0.1μl RNAse inhibitor (40U/μl RRI, TaKaRa)) and immediately stored on dry ice or transferred to a −80°C freezer for long-term storage. All downstream steps were performed as described in Picelli et al 2014 (*19*). Briefly, lysed cells were pre-incubated at 70°C for 3 minutes and stored in ice prior to reverse transcription reaction. RNA was reverse transcribed by adding 5.7μl of RT buffer (2μl 5x RT buffer (SuperScript II, Invitrogen), 0.5μl 100mM DTT, 0.07μl 1M MgCl_2_, 2μl 5M Betaine, 0.25μl RNAse inhibitor (40U/μl RRI, TaKaRa), 0.25μl SuperScript II reverse transcriptase (200U/μl, Invitrogen), 0.2μl TSO (100μM), 0,63μl H_2_O and incubated on a thermocycler for 90 minutes at 42°C, followed by 10 cycles of 50°C and 42°C for 2 minutes each and a 15 minute incubation at 70°C to inactivate the enzyme. The resulting cDNA was amplified for 24 cycles by adding a PCR mix containing IS PCR primers (5’-AAGCAGTGGTATCAACGCAGAGT-3’) (0.25μl, 10μM stock) and KAPA HiFi hotstart ready mix (2× solution, 12.5μl + 2.5μl H2O, Roche, KK2602). PCR conditions were: 98°C/3mins; 24x cycles of (98°C/20s − 67°C/15s − 72°C/6 mins); 72°C/5 mins; 4°C. After PCR was complete, amplified cDNA was washed with AMPure XP beads (Beckman Coulter, A63882) to remove primer dimers and resuspended in nuclease free H_2_O. Sample quality was assessed by running randomly selected samples on a Bioanalyzer (Agilent 2100, High Sensitivity DNA kit, 5067-4626). The concentration of dsDNA in each samples was measured on a Qubit Fluorometric Quantitation device (DNA High Sensitivity Kit, Thermo Fisher Q32851) and samples were stored at −20°C prior to library preparation.

For ***Smart-seq3*** reactions we followed the published protocol as written (https://www.protocols.io/view/smart-seq3-protocol-bcq4ivyw). In brief we sorted labeled T cells into 384 well plates (Armadillo) containing 3μl of Smart-seq3 lysis buffer (0.04μl RNAse inhibitor (40U/μl RRI, TaKaRa), 0.1% Triton X-100, 5% Poly-ethylene Glycol 8000, 0.08μl dNTPs (25mM/each, Thermo Fisher R0182), 0.5μM OligodT30VN (100μM IDT - /5Biosg/ACGAGCATCAGCAGCATACGATTTTTTTTTTTTTTTTTTTTTTTTTTTTTTVN), 2.43μl H_2_O) and were immediately stored at −80°C until reverse transcription step. Prior to adding RT mix the plates were incubated at 72°C on a thermocycler for 10 minutes and stored at 4C immediately until RT mix is added. Reverse transcription was performed by adding 1μL of RT mix (0.1μl Tris-HCl pH 8.3 (1M), 0.12μl NaCl (1M), 0.1μl MgCl2 (100mM), 0.04μl GTP (100mM), 0.32μl DTT (100mM), 0.05μl RNAse Inhibitor (40U/μl RRI, TaKaRa), 0.08μl TSO oligo (100uM, IDT - /5Biosg/AGAGACAGATTGCGCAATGNNNNNNNNrGrGrG), 0.04ul Maxima H-minus RT enzyme (200U/μl), 0.15μl H2O) and incubated on a thermocycler for 90 minutes at 42°C, followed by 10 cycles of 50°C and 42°C for 2 minutes each and a 5 minute incubation at 85°C to inactivate the enzyme. PCR was immediately performed by adding 6ul of PCR mix to each well (2ul Kapa HiFi Hotstart buffer (5x, Roche), dNTPs 0.12ul (25mM/each, Thermo Fisher), MgCl2 (100mM), 0.05ul Fwd Primer (100μM, IDT 5’- TCGTCGGCAGCGTCAGATGTGTATAAGAGACAGATTGCGCAA*T*G-3’), 0.01ul Rev Primer (100μM, IDT – 5’-ACGAGCATCAGCAGCATAC*G*A-3’), 0.2μL DNA Polymerase (1U/μl, Roche), 3.57ul H_2_O). PCR conditions were: 98°C/3mins; 24x cycles of (98°C/20s − 65°C/30s − 72°C/4 mins); 72°C/5 mins; 4°C. For mouse CNS cells 22 cycles was used for preamplifcation. Downstream sample cleanup and quality assessment was performed as described for Smart-seq2. Sample concentrations were measured by incubating 1μl of cDNA from each well with 49μL of a fluorescent dsDNA dye (Quantifluor dsDNA kit, Promega E2670) and measured on a plate reader with fluorescent detectors (504nM Excitation/531nM Emission) and normalized to a standard dilution curve

### Bulk RNA-seq on T cells

For mini-bulk RNA-seq 25 cells were sorted into Smart-seq2 lysis buffer and standard Smart-seq2 reactions were performed with lower numbers of cycles during PCR (20 cycles).

### ATAC-seq

We performed a modified version of the original ATAC-seq protocol optimized for small numbers of input cells (*30*). In brief, we sorted 500-1000 clonally expanded HLA-A2/YFV NS4b-specific CD8+ T cells directly into 22.5μl of ATAC-buffer (12.5μl 2× TD Buffer (Illumina), 0.5μl 1% Digitonin (Promega G9441), 9.5μl H2O) in 96 well plates. After all cells were sorted we added 2.5μl TDE1 enzyme to each well and gently resuspended the solution by pipetting each well 20x careful to avoid adding bubbles. Samples were immediately transferred to a thermocycler set at 37°C and incubated for 30 minutes with the lid set at 50°C. To quench reaction, we added 150μl of ERC Buffer (Qiagen MinElute Reaction Cleanup Kit) to the 25μl ATAC reaction and transferred that 175μl volume to a Qiagen PCR cleanup column containing 150μl of ERC buffer. Samples were centrifuged and washed according to the manufacturer’s protocol and tagmented DNA was eluted in 10μl of H_2_O.

To amplify and index tagmented DNA, we performed a standard PCR using all 10μl of eluted DNA with 25μl 2x NEB High-Fidelity master mix (NEB, MO544), 2.5μl of 25μM forward primer (5’-TCGTCGGCAGCGTCAGATGTGTATAAGAGACAG-3’), 2.5μl of 25μM indexed reverse primers (5’-AATGATACGGCGACCACCGAGATCTACACNNNNNNNNTCGTCGGCAGCGTCAGATGTGTAT-3’) and 10μl H_2_O. PCR conditions were as follows: 72°C/5mins, 98°C/30s; 16x cycles of (98°C/30s − 63°C/30s − 72°C/1min); 4°C hold. Amplified cDNA was size selected using magnetic beads (SPRI, Beckman Coulter B23319) by first incubating with 25ul (into 50μl) beads to remove large cDNA fragments. The unbound liquid was then incubated in a separate well with 50μl (into 75μl) beads to purify the remaining cDNA fragments (size range: approx. 100-800bp). Samples were washed with 80% EtOH and eluted into DNAse/RNAse free H_2_O. Sample quality and concentration were assessed by bioanalyzer (High Sensitivity kit, Agilent) and Qubit (Thermofisher) before being pooled for sequencing.

### Whole Genome Amplification (WGA) of Single Cells

Single CD8+ YFV-specific T cells were index sorted into a 96-well V-bottom plate (Thermofisher) containing 9μl Tris-EDTA (TE) buffer and stored at −20°C for later use. After thawing 1μl of fragmentation buffer (Proteinase K + single cell lysis solution) and cDNA fragments were amplified by PCR according to the manufacturer’s protocol (GenomePlex Single Cell Whole Genome Amplification Kit, Sigma Aldrich WGA4). The amplified libraries were run on a bioanalyzer to assess quality (Agilent) and DNA concentrations were measured on a Qubit Fluorometric Quantitation device (DNA High Sensitivity Kit, Thermo Fisher Q32851) and samples were stored at −20°C prior to library preparation.

### Preparing cDNA libraries for sequencing on Illumina Sequencers

*Smart-seq2 and WGA for CNV analysis*: Libraries were prepared using an in-house tagmentation protocol as previously described (*39*). In brief, 0.5-1ng of cDNA per sample was added to 201l of Tn5 transposase buffer (4μl 40% Poly-ethylene Glycol 8000, 4uL 5x TAPS buffer (50mM TAPS-NaOH, 25mM MgCl_2_ (pH 8.5), 0.1-0.3μl of in-house tn5 transposase, H_2_O to final volume of 20μl – cDNA volume). Samples were incubated on ice and gently pipetted 15-20x to resuspend contents fully. Tn5 binding was carried out on a thermocycler at 55C for 7 minutes and samples were immediately taken after this time and 5μl of SDS (0.2% stock, final concentration 0.02%) was added to quench the Tn5 reaction. Samples were subsequently indexed using Nextera XT 96 dual indexes and KAPA HiFi PCR reagents. 25μl PCR master mix (10ul KaPa HiFi Buffer (5x), 1.5μl dNTP (100mM), 5ul F Primer (N7xx), 5μL R Primer (N5xx), 1μl DNA Polymerase (KaPa HiFi Kit), 2.5μl H2O) was added to the 25μl tagmented cDNA libraries and PCR was performed with the following conditions: 72°C/3mins; 95°C/30s; 10x cycles of (95°C/10s − 55°C/30s − 72°C/30s); 72°C/5 mins; 4°C. Final libraries were cleaned with AMPure XP beads and resuspended in H_2_O. Individual samples were measured and mixed in equimolar ratios for Illumina Sequencing.

*Smart-seq3*: After measuring individual cDNA library concentrations samples were diluted to 100pg/μl followed by a transfer of 100pg cDNA to a new plates for tagmentation. For Smart-seq3 we followed the tagmentation procedures described in the Smart-seq3 protocols.io version 3, with minor modifications. In brief, tagmentation was performed in 1μL of diluted cDNA and 1μL 1x tagmentation mix consisting of 10mM Tris-HCl pH 7.5, 5mM MgCl2, 5% DMF, and 0,1μl ATM (Nextera XT DNA Library Preparation kit, Illumina FC-131-1096), incubated at 55°C for 10min. Tn5 was removed from DNA by addition of 0.5μATM,2% SDS to each well. Following addition of 1,5μl custom illumina indexes (IDT), library amplification PCR was initiated by adding 4μl 1x PCR mix consisting of 1x Phusion Buffer (Thermo Scientific F530L), 0.01 U/µL Phusion DNA polymerase (Thermo Scientific), 0.2 mM dNTP/each (Thermo Scientific). PCR was performed at 3 min 72°C; 30 sec 95°C; 12 cycles of (10 sec 95°C; 30 sec 55°C; 30 sec 72°C); 5 min 72°C in a thermal cycler. After PCR was complete, amplified cDNA was washed with homemade 22% PEG Beads to remove primer dimers and resuspended in nuclease free H_2_O.

*ATAC-seq*: 10μl of tagmented DNA from each sample was used as input (the entire sample) and added to PCR mix for indexing and sample amplification. PCR mix contained (2.5μl primer 1 (25μM Ad1_NoMx), 2.5μl primer 2 (25μM Ad2_xx), 25μL NEB Next HiFi PCR Mix (2x solution, NEB M0544), 10μl H_2_O). PCR conditions were as follows: 72°C/5min; 98°C/30s; 12 x cycles of (98°C/10s – 63°C/30s – 72°C/1min); 4°C. Final libraries were size-selected by performing 2-step bead cleaning (SPRIselect, Beckman Coulter B23318) to remove larger DNA fragments and primer dimers and quality was assessed by Bioanalyzer (Agilent 2100, High Sensitivity DNA kit). Samples were pooled according to indexes for Illumina sequencing.

### Illumina Sequencing

All samples were run by the National Genomics Infrastructure core facility at SciLifeLab in Stockholm, Sweden. For projects: P1902, P3128, P9855, samples were run on an Illumina HiSeq 2500 sequencer using default settings with 2×125 base read length. For Smart-seq3 libraries samples were run on an Illumina NovaSeq 6000 with S4-300 v1.5 flow cells, with 2×150 base read length.

## Data Processing and Analysis

### Data pre-processing for Smart-seq3 of mouse brain cells

For Smart-seq3 data on mouse brain cells, fastq files were generated with bcl2fastq and zUMIs version 2.8.0 or newer was used to process the raw fastq files. Low quality barcodes and UMIs were removed (3 bases < phred 20) before reads were mapped to the mouse genome (mm10) using STAR version 2.7.3. Read counts and error-corrected UMI counts were generated using ensemble gene annotation (GRCm38.91). Cells were filtered as low quality if they did not meet the following criteria; more than 40% of read pairs mapping to exon, at least 20.000 read pairs sequenced, at least 1000 genes detected. The gene expression matrices (UMI counts for introns and exons) for both brains were merged using the merge() function in Seurat v3(*40*). The data were log-normalized with a scale factor of 10000 using the NormalizeData() function followed by linear transformation (scaling) of data. 2000 highly variable features were selected using FindVariableFeatures() followed by PCA and the use of significant PCs (entire dataset: 30; projection neurons: 28, interneurons: 21, oligodendrocytes: 18, astroependymal: 13, immune: 19, vascular: 15) for graph-based clustering (SNN graph calculation and clustering using Louvain). After determining differentially expressed genes, we manually assigned major cell classes to each cluster (Astroependymal, Immune, Neurons, Oligodendrocytes, Vascular) using canonical markers. We then split cells by major cell type, performed subclustering and extensively annotated each cluster based on canonical marker genes from published data and from www.mousebrain.org. At each step, we removed (1) clusters classified with ambiguous labels and (2) outlier cells on the fringes of clusters in UMAP space. We annotated clusters using the same mnemonic identifiers as provided on www.mousebrain.org and added corresponding cell type location and general description as metadata. Finally, we merged all cells into a single file together with metadata and annotations. The filtered cellIDs were exported and used as input for cloneID extraction and clone calling using the TREX Python pipeline^29^. Following clone calling, the obtained cloneIDs were added as metadata to each Seurat object.

### Data pre-processing for Smart-seq2 and Smart-seq3 in Human T-cells

For Smart-seq2 (*19*) data and Smart-seq3 (*10*) data, reads were aligned to the GRCh37 reference with Ensembl version 75 annotations, and raw expression matrices contained 63677 distinct Ensembl gene IDs. Highly variable genes were identified with Scanpy, using the ‘seurat_v3’ method (*40*).

For Smart-seq2 data (single-cell experiments P1902 and P3128), count matrices were normalized to transcripts per million (TPM). Afterwards, genes were filtered out which did not reach a TPM value above 10 for at least 5% of cells in the dataset. For Smart-seq3 data (single-cell experiment YFV2003, *in vivo* donors A,B,C) with unique molecular identifiers (UMI), we filtered out genes which were not expressed (UMI > 0) in at least 5% of cells. UMI data was normalized so that each cell had a total count of 1 million.

All count matrices were then pseudo-log normalized (a count x was normalized to ln(x+1)). T-cell receptor genes were then dropped from the count matrices. For quality control, we inspected the total counts and total number of genes expressed by each cell. We filtered out cells which were outliers in these dimensions, based on visual inspection. Resulting gene expression matrices, together with sample metadata and gene metadata, were saved in AnnData Loom files using the Python package ScanPy (*41*).

### ATAC-seq Peak Calling

ATAC-seq raw sequencing data was analyzed according to https://nf-co.re/atacseq version 1.0.0. The entire pipeline was run with default parameters, except running peak-calling in narrow mode. In summary, after sample quality control and adapter trimming, reads were aligned to the Genome Reference Consortium Human genome build 37 (GRCh37) using bwa. Picard was used to mark duplicate reads and SAMtools/BAMtools for post-filtering of the reads. Normalized bigWig scaled to 1 million mapped reads was created with BEDTools. MACS2 was used for peak calling on the filtered BAM files in narrow-peak mode. A consensus peak set was created with BEDTools, and featureCounts was used to count the reads. All the default parameters, as well as version numbers of the individual tools used in the pipeline can be found at https://nf-co.re/atacseq/1.0.0.

### Identifying TCR sequences for Clonal Analysis

To find clonal populations of T cells we reconstructed TCR*α* and TCR*β* sequences using the software package MIXCR (v3.3)(*42*)

### Copy Number Variation Analysis

Single CD8+ T cell WGA libraries and PBMC unamplified bulk sample libraries subjected to WGS were assessed for quality using FastQC (http://www.bioinformatics.babraham.ac.uk/projects/fastqc/), reads were mapped to the reference genome (human g1k v37 with decoy) using Burrows-Wheeler aligner (BWA-MEM)(*43*). Mapped reads from single cell libraries derived from the same clone were merged into one clonal sample resulting in an average sequencing depth of 0.2x per such clone sample. The unamplified bulk sample was downsampled to 0.2x. The aligned read files were subsequently converted to bed files using bedtools (*44*). Normalized read counts and copy number profiles were obtained by Ginkgo using default parameter settings and a bin size for calling CNV corresponding to 500kbp (*45*).

### Clonally variable genes and controlling FDR, in vivo experiments

To identify clonally variable genes, we used a custom pipeline based on the ANOVA F-statistic (a ratio of variance between groups to variance within groups). The Python implementation of ANOVA F in Scipy (‘f_oneway’) provides lists of p-values for tens of thousands of genes, even with hundreds of cells belonging to dozens of clones, in milliseconds. On the other hand, zero-inflation and other deviations from normality imply that the the p-value obtained from the F-distribution cannot be trusted.

We also utilized the non-parametric Kruskal-Wallis test, in some *in vitro* data, but found that it results largely overlapped with those of the ANOVA F test. In order to identify clonally variable genes, both tests should be used with caution, especially for p-values in the range of 0.01 to 1e-6. We found that genes with ANOVA F-test p-values below 1e-12 were unambiguously clonal, as such p-values did not arise by chance, e.g., when permutation tests were carried out. Our single cell in vitro data sets exhibited many genes which were clonal at the p<1e-12 level, and ANOVA F was sufficient on its own to find large numbers of clonal genes.

For other data sets, we strengthened the ANOVA F-test by performing an approximate permutation test. For this we carried out 1000 permutations of clone labels, to compare the ANOVA F based p-values with a background distribution of p-values for each gene. In fact, we generated 10000 random permutations and took only the 1000 permutations which most thoroughly scrambled the clone labels. E.g., a permutation which sends labels AAABBBCCC to BBBAAACCC would receive a minimal “scrambling score,” since it has no real effect on the clonal groups. More precisely, the scrambling score of a permutation was defined as the sum of the numbers of unique clone labels received within each real clone group. For example, a permutation sending real labels AAABBBCCC to ABCBBACCA would receive a score of 3+2+2=7, since the previous single-label groups AAA, BBB, CCC received 3 distinct labels (ABC) and 2 distinct labels (BBA) and 2 distinct labels (CCA), respectively.

This permutation procedure enabled us to estimate the excess of clonally variable genes, defined as those with p<0.05, for *in vivo* experiments. To estimate this excess, we compared the number of clonally variable genes found with real clone labels to the median and 95th percentile among 1000 scrambled clone labels. Furthermore, by choosing more stringent p-value cutoffs, we were able to identify smaller sets of clonally variable genes while controlling the false discovery rate, e.g., finding 50 clonally variable genes with an estimated 1 false discovery. To estimate the number of false discoveries, we considered an additional 100 permutations of clone labels -- since the previous 1000 permutations were used in choosing an appropriate p-value cutoff.

### Differentiation state ranking by PC1

To rank the differentiation state for in vivo cells, we began by merging the gene expression matrices for three donors (A,B,C), taking those genes which were clonally variable in at least one donor and expressed by all donors. This led to a set of 107 genes, on which we performed principal component analysis.

As expected, the largest loadings in PC1 were genes associated with differentiation state, like *GZMH*, *SELL*, etc. Genes were annotated as differentiation markers if they had a loading of above 0.1 or below −0.1, in PC1. We assigned each cell a score, between −0.5 and 0.5, based on PC1, according to the following procedure: the PC1 range was split into two equal-length bins, and then linearly normalized to take values between −0.5 and 0.5, with the bin-divider at zero. The result was then negated, if necessary, so that *GZMH* (a marker of highly differentiated T cells) was positively correlated with the normalized PC1 value. The resulting number was used as a score for differentiation state, with −0.5 indicating a less differentiated state and 0.5 a more highly differentiated state for each cell.

### Machine learning for clonal gene expression signatures

A machine-learning classifier called a linear support vector machine (linear SVM, or simply SVM) was applied to single-cell gene expression matrices, to determine whether clonality could be predicted from gene expression in a supervised setting. The SVM pipeline comprised three steps: a min/max scaler to scale all gene expression (previously pseudolog-normalized) to a common interval, the selection of k most clonal genes based on ANOVA F statistic, and then the application of the linear SVM with a misclassification penalty parameter C. The linear SVM classifier aims to separate each clone from all of the others (one v. all method), using a weighted combination of the k selected clonal genes (a “metagene”) as an SVM hyperplane.

The two hyperparameters for this pipeline are the number of genes used for prediction (k), and the penalty parameter (C). Grid-search with 5-fold cross-validation was applied to find hyperparameters which optimized the predictive accuracy of the SVM. This search initially considered between 2 and 300 genes, and loss penalties C from 0.001 to 100.0. The C parameter had little effect, once it was at least 0.1, and so we focused on a more refined analysis of the number of genes on predictive accuracy. The same pipeline was applied with C=0.1, and between 1 and 800 genes, to record the accuracy of clonality prediction. The entire process was repeated with shuffled clone labels, in order to find an expected level of accuracy under a null hypothesis. All machine learning pipelines were implemented in Python, using the scikit-learn package (*46*).

### Predicting clonality and confusion matrices

To understand whether predictive accuracy was greater or less for specific clones, we repeatedly (100 times) ran the SVM pipeline with optimal parameters k,C, to see how often cells from one clone were (correctly or incorrectly) classified as belonging to another clone based on gene expression. The results were recorded in “confusion matrices” whose diagonal reflects the proportions of each clone that were correctly classified. For these confusion matrices, 80% of cells were used to train the SVM and 20% of cells were then held aside for testing predictive accuracy, to match the 5-fold cross-validation earlier. Confusion matrices were also produced with fewer training cells (67% and 50%), but the resulting accuracy of clonal prediction did not greatly suffer.

This method was adapted to study cross-well clonal prediction in the P3128 dataset (Fig. 3). In this case, cells from four clones were distributed into eight wells. Rather than randomly splitting the cells 80/20 into training/testing sets, cells from four wells were used for training the SVM. Following training, the SVM was used to predict the clonality of cells from the remaining four wells. One well contained a mix of two clones and was therefore unsuitable for training the SVM to predict clonality. The cells from that well were therefore held out in the testing set in all cases. This was repeated 100 times, switching wells used for training with those used for testing each time, using penalty parameter C=1.0 and k=200 (200 genes).

### ANOVA and Nested ANOVA for clonal and well-significant genes

Excluding the odd mixed-clone well, the cells from sister clones (103 cells from clones 1,11,13,54) in P3128 belonged to 4 clones, which were then split among 8 wells. In order to identify long-term clonally significant genes, and distinguish them from potentially short-term well-dependent genes, we applied a nested ANOVA design. This first step applies a standard ANOVA F test to measure the clonal significance of each gene. After this, the nested ANOVA looks for significant differences between the two wells within each clone, applying the equivalent of a t-test (ANOVA F for two wells). Results of the nested ANOVA are reported as unadjusted p-values.

### Data preprocessing for bulk Smart-seq2 and ATAC-seq

In experiment P9855, we gathered gene expression information for 70 samples, each with 25 bulks. These 70 samples came from 24 clones based on TCR. Preprocessing for these bulks followed the same pipeline as single-cell Smart-seq2, including identification of highly variable genes, TPM-normalization, pseudo-log normalization, dropping TCR genes, and examination of total counts and genes for quality control. After quality control, we kept 48 samples, representing three 25-cell mini-bulks from each of 16 clones. Clonal genes were assessed by ANOVA F statistic as before.

ATAC data was obtained for 29 bulks, consisting of between 269 and 1000 cells (with most bulks having 1000 cells). These include 6 pairs of biological replicates (clones 1,15,22,23,8,9) and one pair of sister clones (5a, 5b) sharing TCR sequence. ATAC data contained the heights of 80599 peaks for each sample, further annotated with genomic location, and type (intron, promoter-TSS, etc.). Peaks were removed that were annotated as promoters for TCR genes, and also if they were located within 100 Kbps upstream or downstream of the TSS for a TCR gene. This removed 749 peaks. Peaks were filtered to exclude those peaks which never rose above a height of 30. This removed about two thirds of the peaks, leaving 26040 peaks. The ATAC peak height matrix was psuedolog-normalized, then stored, with all annotations, in an AnnData Loom file, using the Python package ScanPy (*41*).

### Dimensional reduction and measuring similarity after ATAC-seq

With only 29 samples and 26,040 peak heights, Euclidean distance is not expected to adequately convey the similarities among samples. Therefore, we considered the distance between samples after dimensional reduction by principal component analysis (PCA with 1-20 PCs). After dimensional reduction, we compared pairwise distances between (1) replicate pairs, (2) the sister clones 5a/5b, and (3) non-replicate samples.

Using only PCA with at least 5 principal components, we found that pairwise distances between replicates were 5% of the pairwise distances between non-replicate samples, on average. The distance between sister clones was greater, but still below 50% of the distance between non-replicate samples. While this level of proximity is not particularly significant in 1 or 2 dimensions, it is very significant (beyond two standard deviations below the mean) when one gets to 15 PCs or more. This reflects a general fact about high-dimensional data – Euclidean pairwise distances naturally grow larger as one adds more dimensions, but the standard deviation among these pairwise distances remains stable. For example, if one chooses uniformly random points in a d-dimensional box (with coordinates between 0 and 1), then the expected pairwise distance grows proportionally to the square root of d. The standard deviation among these distances remains constant. For normal distributions, the same is true as d grows large. Thus, in high dimensions, it becomes much rarer to see points that are – for example – half as far apart as a randomly chosen pair of points.

### Variability of ATAC peaks using biological replicates

The six pairs of biological replicates enabled us to analyze technical noise in ATAC peak heights. When analyzing the 6 replicate pairs (12 samples), it became evident that the mean-variance relationship was complicated, especially for lower peaks. Thus, we applied a non-parametric approach with smoothing in order to model the dependence of technical noise on peak height.

We first pooled all 6 replicate pairs among 26040 peak intervals to obtain 156,240 replicate-peak-pairs (RPPs). Each RPP therefore comprised a peak of lowest height (*h_i_*) and tallest height (*H_i_*). If one considers another peak height (*h*), the tallest height one might expect among a replicate would be max{*H_i_ : h_i_ ≤ h* }; in other words, the largest height among replicate pairs whose low peak is lower than *h*. To deal with unexpected noise for peaks near zero, we conservatively shifted this to max{ *H_i_: h_i_ ≤ h* + 10 }. Based on this, we defined the maximum expected gap (*MEG(h)*) for a height *h* to be

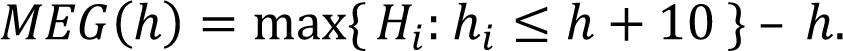

By nature, this function exhibited frequent discontinuities, and so we applied a cubic smoothing filter (Savitzky-Golay) with large window (window-size 601 among data between 0 and 700). This gave a function *MEG_sm_(h)* which can be summarized as the maximum expected technical noise, for a peak interval whose lowest occurring height is *h*.

We used this expectation of technical noise to normalize a metric of peak height variation among all peak intervals (CREs). Namely, for any such peak interval, there is a lowest height *h* and highest height *H*, among the 29 samples. The ‘gap’ of the peak interval is just the difference *H – h*, and we defined the ‘relative peak variability’ to be the gap normalized by the maximum expected technical noise:

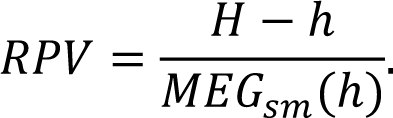

### Correlation between of genes and nearby peaks

When considering ATAC peak intervals “near” a gene, we considered all peaks whose midpoint was within the length of the gene or 50kbp upstream of the gene. We added another 1000bp of tolerance to avoid close misses, narrow peaks, etc., to create a window around each gene.

To correlate gene expression and peak height, we took clonal averages of each -- averaging the three 25-cell samples for each of the 16 clones within the gene expression matrix and averaging the replicate samples to find ATAC peak height averages for the same 16 clones. Pearson correlation coefficients were used throughout.

### Principal component regression and covariant peaks

For some genes, we found numerous nearby CREs whose ATAC peak height was highly correlated with gene expression. To assess the *independent* contributions of nearby CREs, we performed principal component regression (*47*). For each gene, we considered all CREs within the usual window, restricting to those that reached a height of 30 as before. Among these peaks, we restricted to those whose correlation with gene expression reached a threshold of *R*^2^ > 0.05, a light supervision to remove some peaks which were irrelevant to gene expression.

We computed the correlation matrix of the remaining peaks and defined ‘eigenpeaks’ to be the eigenvectors of this correlation matrix, i.e., the eigenpeaks for a given gene are the principal components of the relevant nearby peaks. By construction, eigenpeaks do not ‘see’ gene expression (except for the light initial filter) and eigenpeaks have zero correlation with each other. Subsequently, we computed the correlation of the eigenpeaks with gene expression. Eigenpeaks which are highly correlated to gene expression reflect additive combinations of CREs that predict gene expression. Even when there were many (up to nine) highly correlated peaks near a gene, there was rarely more than one highly correlated eigenpeak (**Fig. S7, D and E**). This indicates that nearby CREs typically act in concert to regulate gene expression.

## Data Availability

All sequencing data will be deposited and made available to the scientific community upon request pending publication.

## Code Availability

Processed data and code needed to generate figures from this study are available online at GitHub at: https://github.com/MartyWeissman/ClonalOmics and contains python notebooks with instructions for data processing as well as all data necessary to run notebooks (processed sequencing data, metadata files).

